# Heterogeneous substrate binding in a glutamate transporter homologue

**DOI:** 10.1101/2021.07.08.451670

**Authors:** Krishna D. Reddy, Didar Ciftci, Amanda Scopelliti, Olga Boudker

## Abstract

Integral membrane glutamate transporters couple the concentrative substrate transport to ion gradients. There is a wealth of structural and mechanistic information about this family. Recent studies revealed transport rate heterogeneity in an archaeal homologue Glt_Ph_, inconsistent with simple kinetic models, but its structural and mechanistic determinants remain undefined. In a mutant Glt_Ph_, which exclusively populates the outward-facing state, we demonstrate the co-existence of at least two sub-states in slow equilibrium binding the substrate with different apparent affinities. Wild-type Glt_Ph_ shows similar binding properties, and modulation of the sub-state equilibrium correlates with transport rates. Following binding, the low-affinity sub-state of the mutant is transient. Consistently, cryo-EM on samples frozen within seconds after substrate addition reveals the presence of structural classes with perturbed helical packing of the extracellular half of the transport domain in regions adjacent to the binding site. In contrast, an equilibrated structure does not show such classes. The structure at 2.2 Å resolution details a pattern of waters in the intracellular half of the domain and resolves classes with subtle differences in the substrate-binding site. We hypothesize that the rigid cytoplasmic half of the domain mediates substrate and ion recognition and coupling, while the extracellular labile half sets the affinity and dynamic properties.

## MAIN TEXT

Membrane glutamate transporters pump substrates against their concentration gradients, serving critical biological functions across all kingdoms of life. In mammals, excitatory amino acid transporters (EAATs) recycle glutamate from the synaptic cleft into the glia (Freidman et al., 2020). In prokaryotes, orthologs uptake nutrients, including glutamate, aspartate, neutral amino acids, or dicarboxylic acids (Burguiere, Auger, Hullo, Danchin, & Martin-Verstraete, 2004; Kim et al., 2002; Youn, Jolkver, Kramer, Marin, & Wendisch, 2009). Transporters utilize energy from downhill ionic electrochemical gradients to carry concentrative substrate uptake. EAATs rely on inward Na^+^ and proton gradients and an outward K^+^ gradient (Zerangue & Kavanaugh, 1996). Prokaryotes couple transport to either proton or Na^+^gradients (Ryan, Compton, & Mindell, 2009; Tolner, Ubbink-Kok, Poolman, & Konings, 1995).

These transporters are homotrimers with each protomer composed of a rigid scaffold trimerization domain, and a mobile transport domain containing the ligand-binding sites. Protomers functions independently (Erkens, Hanelt, Goudsmits, Slotboom, & van Oijen, 2013; Georgieva, Borbat, Ginter, Freed, & Boudker, 2013; Grewer et al., 2005; Koch, Brown, & Larsson, 2007; Koch & Larsson, 2005; Riederer et al., 2018; Ruan et al., 2017). Transport domains translocate ligands across membranes by moving ~15 Å from the outward-facing state (OFS) to the inward-facing state (IFS), in an ‘elevator’ motion (Arkhipova, Guskov, & Slotboom, 2020; Garaeva, Guskov, Slotboom, & Paulino, 2019; Qiu, Matthies, Fortea, Yu, & Boudker, 2020; Reyes, Ginter, & Boudker, 2009). Studies in archaeal Na^+^-coupled transporters Glt_Ph_ and Glt_Tk_ led to a simple kinetic model of transport (Alleva et al., 2020; Arkhipova et al., 2020; Boudker, Ryan, Yernool, Shimamoto, & Gouaux, 2007; Guskov, Jensen, Faustino, Marrink, & Slotboom, 2016; Oh & Boudker, 2018; Reyes, Oh, & Boudker, 2013; Riederer & Valiyaveetil, 2019; Verdon, Oh, Serio, & Boudker, 2014; X. Wang & Boudker, 2020). Briefly, in the OFS, ion binding to Na1 and Na3 sites reveals the substrate- and an additional sodium (Na2) binding site through an opening of helical hairpin 2 (HP2), also preventing the translocation of Na^+^-only bound transport domain. Subsequent binding of the substrate and Na2 closes HP2, allowing translocation to the IFS and ligand release into the cytoplasm.

Recently, high-speed atomic force microscopy, single-molecule Förster resonance energy transfer (smFRET) total internal reflection fluorescence (TIRF) microscopy, and ^19^F-NMR revealed a more complex picture of Glt_Ph_ transport and dynamics (Akyuz, Altman, Blanchard, & Boudker, 2013; Akyuz et al., 2015; Ciftci et al., 2020; Ciftci et al., 2021; Erkens et al., 2013; Huang et al., 2020; Huysmans, Ciftci, Wang, Blanchard, & Boudker, 2021; Matin, Heath, Huysmans, Boudker, & Scheuring, 2020).

These studies established the existence of additional conformational sub-states in OFS and IFS, of which some translocate and transport at different rates. Though it is expected that cryo-EM studies would resolve these proposed conformational sub-states from heterogeneous datasets, this so far does not appear to be the case (Arkhipova et al., 2020; X. Wang & Boudker, 2020). Therefore, the structural and mechanistic determinants of kinetic heterogeneity remain unclear.

Using a Glt_Ph_ mutant that exclusively occupies the OFS, we show that substrate binding in the OFS is heterogeneous, consistent with at least two sub-states with different affinities. Salt composition and temperature modulate the sub-states populations, suggesting that they are in equilibrium. However, the conformational exchange must be slow, on the order of at least tens of seconds, for the sub-states to manifest in binding isotherms. Notably, a similar conformational equilibrium also exists in the wild-type (WT) protein and potentially contributes to heterogeneous transport kinetics. The substrate-bound high-affinity state of the mutant transporter is expected to predominate after equilibration. Thus, to gain insights into the structure of the transient low-affinity sub-state, we conducted an extensive analysis of the cryo-EM imaging data collected on the transporter frozen within seconds after substrate addition. The identified structural classes reveal subtle differences. Specifically, we observed a subset of structural classes with differently packed helices in the extracellular half of the transport domain adjacent to the binding site, suggesting that the region is labile. We hypothesize that the ensemble of transient, more dynamic, less uniquely packed conformations comprises the low-affinity sub-state. Images of an equilibrated protein produced reconstructions at a uniquely high 2.2 Å resolution, revealing a complement of structured waters in the cytoplasmic side of the transport domain that may contribute to its conformational rigidity. We did not observe a conformational heterogeneity of the extracellular half of the transport domain in these data, which we attribute to the relaxation of the protein to the higher affinity state. Classifications instead revealed subtle differences in the substrate-binding site and the global orientations of the transport domains, which could also contribute to kinetic heterogeneity. Our results provide a framework where modal kinetic behavior demonstrated by Glt_Ph_, and other proteins may be a result of subtle but long-lived structural differences.

## RESULTS

### P-Glt_Ph_ (S279E/D405N) reveals two outward-facing substrate-binding conformations modulated by temperature and salts

We generated a mutant, P-Glt_Ph_, that eliminates Na^+^ binding to Na1 (D405N) and introduces a protonatable residue at the tip of HP1 (S279E), mimicking amino acid sequence features of proton-coupled orthologues (**Figure 1** - **Supplementary Figure 1a-b**). Our original intention was to test the pH dependence of this mutant. While proton gradients stimulated P-Glt_Ph_ activity in the presence of Na^+^ (**Figure 1 - Supplementary Figure 1c**), this is not the focus of this study. We observed that P-Glt_Ph_ had greatly diminished transport compared to the WT transporter (Ryan et al., 2009), prompting us to test its ability to translocate substrate using smFRET. We introduced a single cysteine mutation into a cysteine-free background (P-Glt_Ph_ C321A/N378C), purified the protein in DDM, labeled with donor and acceptor fluorophores, and analyzed by smFRET as in earlier studies (Akyuz et al., 2013; Akyuz et al., 2015; Huysmans et al., 2021). Conformations of protomer pairs within individual Glt_Ph_ trimers can be distinguished by FRET efficiency (*E_FRET_*) as either both in OFS (~0.4), a mixture of OFS and IFS (~0.6), or both in IFS (~0.8). Most P-Glt_Ph_ molecules occupy a low *E_FRET_* state in the presence of 500 mM sodium salts, regardless of anion or substrate presence (**Figure 1a-b**). Thus, this mutant is mainly in OFS or the intermediate iOFS (Huang et al., 2020; Verdon & Boudker, 2012), which our smFRET setup cannot distinguish.

**Figure 1.**
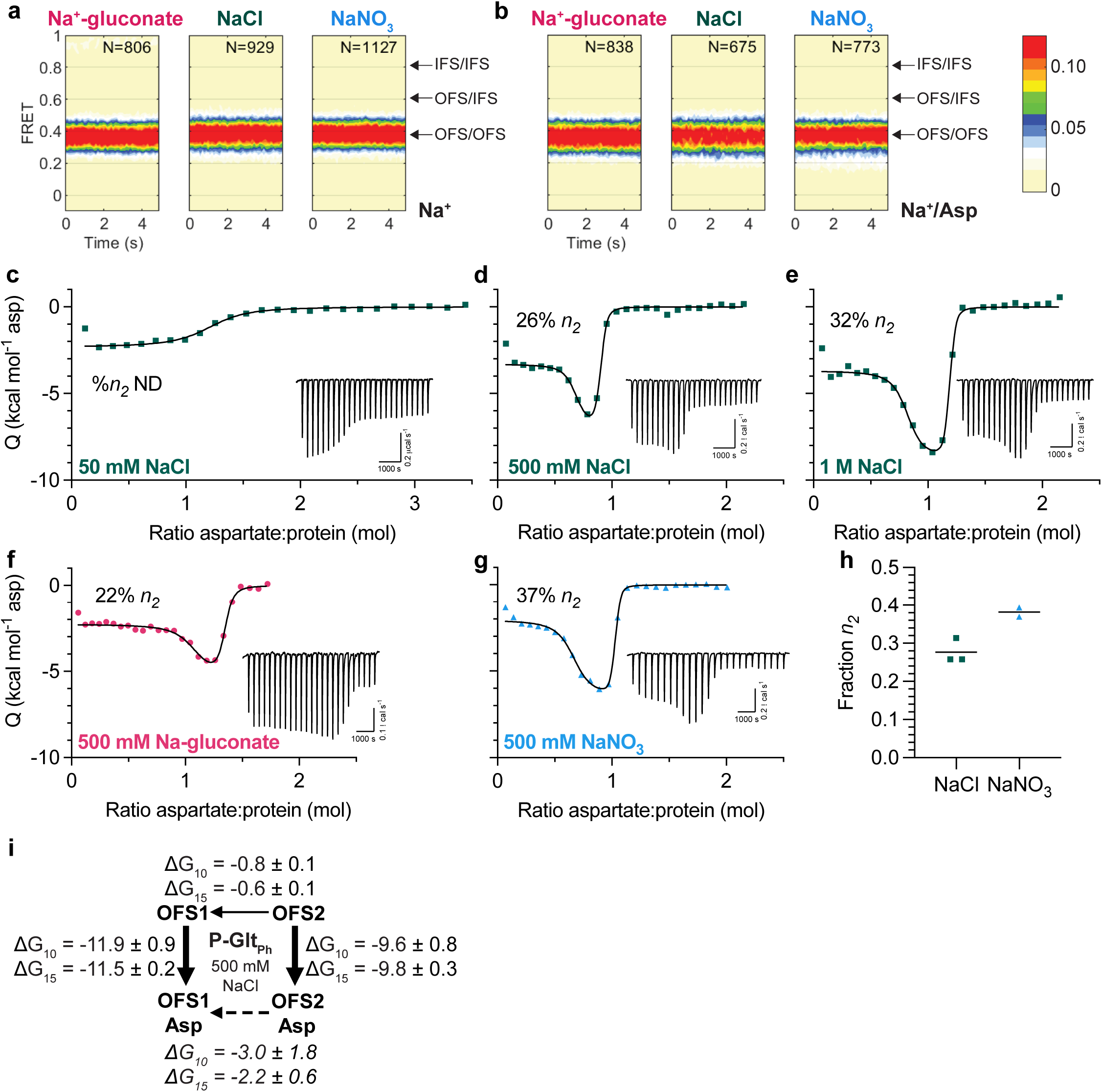
Two outward-facing substrate-binding states in P-Glt_Ph_ (S279E/D405N). (a-b) FRET efficiency population histograms of P-Glt_Ph_ in the presence of 500 mM sodium salts, either in the absence (a) or presence (b) of 1 mM L-Asp. *N* is the number of molecules analyzed. Data shown are an aggregate of two independent experiments. Population contour plots are color-coded from tan (lowest population) to red (highest). Expected conformations according to *E_FRET_* values are indicated by arrows. (c-f) ITC experimients on P-Glt_Ph_ at 15°C were performied at least twice on independently prepared protein samples with similar results. Insets show the thermal power with the corresponding scales. (c-e) Aspartate binding isotherms derived from the ITC experiments in the presence of different amounts of NaCl (green squares): 50 mM (c); 500 mM (d); and 1 M (e). The 50 mM data were fitted to the single-state model, with fitted *K_d_* = 917 nM, Δ*H* = −2.3 kcal mol^−1^, and an apparent number of binding sites, *n* = 1.18. 500 mM NaCl and 1 M NaCl data were fitted to the two-state binding model. The 500 mM NaCl data gave the following fitted parameters for the two states: *K*_d_: 1.3 and 60.8 nM, Δ*H*: −3.3 and −7.1 kcal mol^−1^, *n*: 0.64 and 0.22. The 1M NaCl data: *K_d_*: 0.5 and 27.4 nM, Δ*H*: −3.7 and −8.7 kcal mol^−1^, *n*: 0.77 and 0.38. (g-h) Aspartate binding isotherms were obtained in 500 mM Na-gluconate (f, red circles), or NaNO_3_ (g, blue triangles). All data were fitted to the two-state model. The 500 mM Na-gluconate data: *K*_d_: 4.4 and 138.3 nM, Δ*H*: −2.3 and −5.5 kcal mol^−1^, *n*: 1.00 and 0.28. The 500 mM NaNO_3_ data: *K*_d_: 0.9 and 34.3 nM, Δ*H*: −2.1 and −6.5 kcal mol^−1^, *n*: 0.62 and 0.37. (h) Comparison of the *n_2_* fraction in 500 mM NaCl or NaNO_3_. Each point is an independent experiment. (i) Schematic representations of the conformational and binding equilibria obtained experimentally at 10°C and 15°C in 500 mM NaCl (solid lines) or inferred (dashed lines). The thermodynamic parameters were estimated under the assumptions that there are two non-exchanging binding states. Directions of the arrows indicate directions of the free energy changes, *ΔG-s*, shown. All values are in kcal mol^−1^. Binding *ΔG*-s are from Supplementary Tables 1 and 2. The free energy differences between sodium-bound outward-facing states were calculated from equilibrium constants *K_eq_*= *n_1_*/*n_2_*. Thin lines represent steps that are slow on the time scale of ITC experiments.

Further characterization of substrate binding to the mutant yielded results incompatible with our current understanding of the glutamate transporter mechanism. We expected to observe simple 1:1 binding of aspartate to a single P-Glt_Ph_ binding site in isothermal titration calorimetry (ITC) experiments. Instead, we observed bimodal binding isotherms in 500 mM NaCl at 15°C (**Figure 1d**). Because this mutant exclusively occupies OFS, this result suggests heterogeneity in substrate binding to this state. A ‘two-state’ model assuming two independent, non-identical binding states (OFS1 and OFS2) (Freire, Schon, & Velazquez-Campoy, 2009) is the simplest model that fits this data reliably. Several other binding models for complex equilibria, including cooperative and sequential binding, cannot fit our data. Notably, lack of coupling between protomers has been well-established (Erkens et al., 2013; Georgieva et al., 2013; Riederer et al., 2018; Ruan et al., 2017). Taken together, the data suggest there are two dominant binding states, though there could be additional underlying complexity. Furthermore, it is likely that these two states represent two different conformations of the same site in the transporter population rather than two distinct binding sites within a protomer, since the sums of the apparent stoichiometries (*n_1_* and *n_2_* values) averaged 0.97 ± 0.26 (range of 0.74-1.53, *N*=6) (**Figure 1 - Supplementary Tables 1 and 2**). The bimodal isotherms reflect the presence of a conformation with a higher affinity and lower exothermic binding enthalpy (OFS1) and a conformation with a lower affinity and higher exothermic enthalpy (OFS2) (Brautigam, 2015; Le, Buscaglia, Chaires, & Lewis, 2013). The two conformations must interconvert only slowly, or not at all, during the ITC experiment to manifest two distinct binding states. Notably, we do not observe bimodal isotherms in 50 mM NaCl (**Figure 1c**), likely because the lower substrate affinity at lower Na^+^ concentrations (Boudker et al., 2007; Reyes et al., 2013) blurs the distinctions between the states.

Increasing NaCl concentration to 1 M led to qualitative differences in the isotherms, attributed to the increased OFS2 fraction and its exothermic enthalpy (**Figure 1e**). The measured Na^+^*K_D_-s* for WT Glt_Ph_, 99-170 mM (Hanelt, Jensen, Wunnicke, & Slotboom, 2015; Reyes et al., 2013; Riederer & Valiyaveetil, 2019), suggest that Na3 site is saturated at 500 mM NaCl. Thus, the OFS2 population is unlikely to depend on the extent of Na^+^ binding and instead might reflect general salting effects. To test this, we determined OFS2 fraction in the presence of 500 mM sodium salts containing anions on different ends of the Hofmeister lyotropic series (gluconate^−^ < Cl^−^ < NO3^−^) (Zhang & Cremer, 2006). Gluconate and nitrate have the opposite effects on protein structure; respectively, they decrease and increase the solubility of nonpolar moieties - “salting out” and “salting in” effects. The biphasic shape of the binding isotherms is less pronounced in gluconate than chloride and nitrate (**Figure 1d, f-g**). Fitted OFS2 fraction increases from ~28 % in NaCl to ~38 % in NaNO3 (**Figure 1h**). It decreases in Na^+^-gluconate, though the binding parameters were difficult to model reproducibly. Decreasing temperature to 10°C resulted in the OFS2 fraction falling to ~20 % in NaCl (**Figure 1 - Supplementary Tables 1 and 2**). Increasing temperatures above 15 °C resulted in protein aggregation.

The observation that temperature and chaotropic salts increase the OFS2 fraction suggests that OFS1 and OFS2 are in a slow equilibrium. Furthermore, OFS2 likely features increased solvent accessibility of hydrophobic regions. We estimated the free energy differences between OFS1 and OFS2 based on the measured populations and binding free energies at 10 and 15°C under the assumption that the states do not exchange significantly during the ITC experiment (**Figure 1i**). OFS1 and OFS2 are nearly isoenergetic before substrate binding, but OFS1 predominates when bound to L-Asp, reflecting higher affinity. Transiency of OFS2 may explain why alternate substrate-binding conformations have not been visible in structures. Nevertheless, it must be kinetically stable over the course of ITC titrations. Notably, nanomolar apparent binding affinities measured for P-Glt_Ph_ suggest that substrate dissociation contributes little to the binding process observed in ITC. Thus, the two states may differ primarily in the binding on-rates, with the “high-affinity” state binding substrate faster than the “low-affinity” state. Regardless, these results strongly suggest that there is conformational heterogeneity in the transporter, manifesting in different binding mechanisms.

### Heterogeneous substrate binding in wild-type Glt_Ph_

We also performed ITC experiments on the WT protein. When we used high protein concentrations to increase experimental sensitivity, we observed unusual features in binding isotherms (**Figure 2a-c**). Specifically, the initial L-Asp injections do not have constant heats as expected for a high-affinity single-site binding process. Instead, injection heats steadily decrease until an abrupt drop occurs when the ligand saturates the protein. The two-state model, where the two affinities are close but not identical and the higher-affinity state has a higher exothermic binding enthalpy, fits data well, though the binding parameters are not uniquely determined (**Figure 2 - Supplementary Figure 1a**). Like in P-Glt_Ph_, we observed qualitative differences in the isotherms in different salts and temperatures. The isotherm in 500 mM Na-gluconate at 15°C has a particularly unusual shape (**Figure 2a**) but becomes more reminiscent of a single-site binding in more chaotropic salts (**Figure 2b-c**) or at higher temperatures (**Figure 2d-e**). As in P-Glt_Ph_, changes in the state populations can account for the changing isotherm shapes (**Figure 2 - Supplementary Figure 1b**), though faster exchange between states, or altered enthalpies could also contribute. Collectively, our data suggest that wild-type Glt_Ph_ has multiple binding states in temperature- and salt-modulated equilibrium.

**Figure 2.**
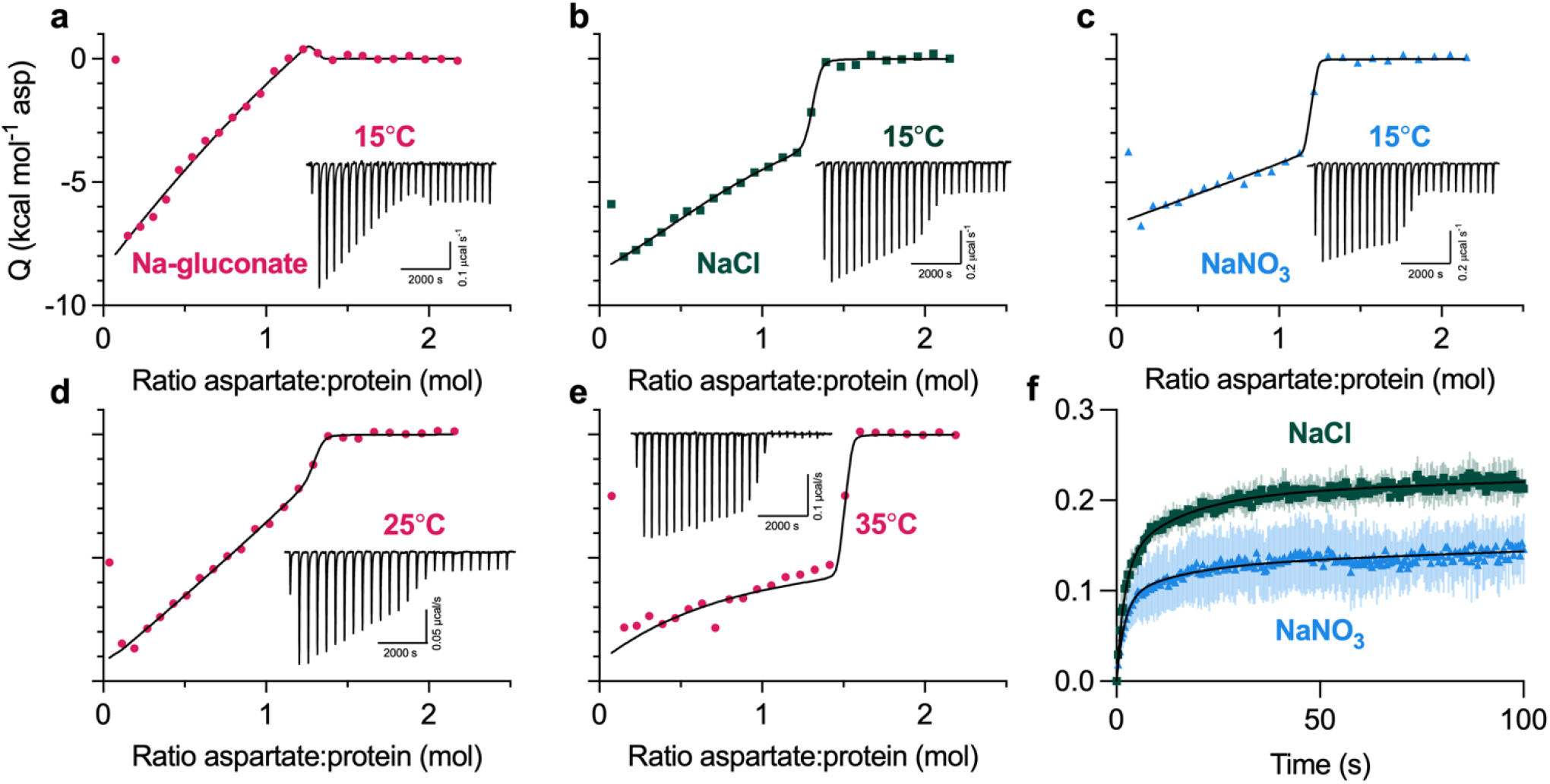
Heterogeneous substrate binding in WT Glt_Ph_. (a-c) Aspartate binding isotherms derived from the ITC experiments performed at 15°C in the presence of 500 mM (a) Na-gluconate (red circles); (b) NaCl (green squares); or (c) NaNO_3_ (blue triangles). (d-e) Aspartate binding isotherms in 500 mM Na-gluconate at 25°C (d) or 35°C (e). Experiments in (a-e) were performed at least twice on independently prepared protein samples, producing similar results. All data were fitted to the two-state model (black lines); however, the binding parameters are not uniquely determined (see Supplementary Figure 2a for further information). Insets show the thermal power with the corresponding scales. (f) Aspartate transport of Glt_Ph_ (C321A/N378C) was measured using the single-transporter FRET-based assay in NaNO_3_ (blue) or NaCl (green). Transport was initiated by perfusing surface-immobilized proteo-liposomes with buffer containing 200 mM sodium salt, 1 μM valinomycin, and 1 μM L-Asp. Data are shown as fractions of total observable transport over time, fitted to tri-exponential functions (black lines). The fitted parameters are in Figure 2 – Supplementary Figure 2b. Data are means and SE from three independent experiments.

WT transporter in saturating Na^+^ concentrations is predominantly in the OFS (Akyuz et al., 2013; Akyuz et al., 2015), but we cannot exclude that inward-facing protomers contribute to heterogeneous L-Asp binding. However, binding of the transport blocker TFB-TBOA, which has a 100-fold preference for OFS (Boudker et al., 2007; McIlwain, Vandenberg, & Ryan, 2016; X. Wang & Boudker, 2020), also produced bimodal ITC isotherms reminiscent of P-Glt_Ph_ (**Figure 2 – Supplementary Figure 2a-c**). Due to the lower TFB-TBOA affinity, we could not determine precise binding parameters and quantify the salt effects. When Glt_Ph_ saturated with TFB-TBOA was competed with L-Asp, we again observed bimodal isotherms (**Figure 2 – Supplementary Figure 2d-f**). Thus, the inhibitor binding is heterogeneous, and some level of conformational heterogeneity persists after binding.

We used a recently developed smFRET-based single-transporter assay to test if salt-modulated state populations correlated with transport rates (Ciftci et al., 2020). P-Glt_Ph_ C321A/N378C mutant was labeled with PEG_11_-biotin and N-ethyl maleimide (NEM) and reconstituted into liposomes; low protein-to-lipid ratios enriched vesicles containing at most one Glt_Ph_ trimer. The proteoliposomes were then loaded with periplasmic glutamate/aspartate binding protein (PEB1a) Y198F/N73C/K149C mutant with reduced aspartate affinity labeled with maleimide-activated donor (LD555P) and acceptor (LD655) fluorophores (referred to altogether as ccPEB1a-Y198F). The proteoliposomes were immobilized in microscope chambers via biotinylated transporter and assayed for transport following perfusion with saturating Na^+^ and L-Asp concentrations. An increase in mean *E_FRET_* from ~0.6 to ~0.8 reflects saturation of the ccPEB1a-Y198F sensor by L-Asp molecules transported into vesicles. This assay previously established kinetic heterogeneity in WT Glt_Ph_ transport (Ciftci et al., 2020), where at least three observable populations (“fast,” “intermediate,” and “slow”) transport at vastly different rates and all contribute to mean uptake measured in bulk. Most WT transporters are “slow,” with turnover times of tens to hundreds of seconds.

Because Glt_Ph_ mediates an uncoupled anion conductance, which dissipates the buildup of membrane potential due to electrogenic transport, it shows faster uptake in the presence of more permeant anions (gluconate^−^ < Cl^−^ < NO_3_^−^) (Ryan & Mindell, 2007). Thus, we measured transport in K^+^-loaded proteoliposomes in the presence of ionophore valinomycin clamping the potential. Tri-exponential fits of the uptake kinetics suggest that most molecules are in the “slow” transporting population regardless of the salt, as expected. The fractions of the “slow” transporters were similar in Na-gluconate (81.1 ± 3.1) and NaCl (79.5 ± 3.4 %) conditions but increased in NaNO_3_ conditions (87.3 ± 2.2 %) (**Figure 2f and Figure 2 – Supplementary Figure 3a-b**). Therefore NaNO_3_-favored OFS2 conformation might correlate with a slower transporter population.

### Transient transport domain structures following substrate binding

smFRET has shown that P-Glt_Ph_ is exclusively outward-facing, making this mutant an excellent model to dissect differences between OFS1 and OFS2 using cryo-EM, where structural heterogeneity should reflect the binding heterogeneity. Because OFS2 is transient and OFS1 predominates at equilibrium, we optimized conditions to increase the probability of imaging OFS2. ITC analysis showed that elevated temperatures and chaotropic salts increased the OFS2 fraction (**Figure 1**). Thus, we pre-equilibrated P-Glt_Ph_ in 250 mM NaNO_3_ at 25°C and froze grids within ~5 s after adding 1mM L-Asp (Dataset A).

When we refined particles with imposed C3 symmetry, we obtained density maps with an overall resolution of 3.0 Å (**Figure 3 – Supplementary Figure 1**). To maximally retain heterogeneity, we used a data processing approach designed to pick the highest-quality particles regardless of conformation (Su et al., 2020). We then used symmetry expansion and focused classification of single protomers into ten classes followed by local refinement, a processing approach that previously revealed OFS and iOFS in the WT Glt_Ph_ ensemble (Huang et al., 2020). We did not find any iOFS classes and only observed OFS classes with similar overall structures. We refined models for four classes with the highest resolution, from 3.15 to 3.85 Å (**Figure 3 – Supplementary Figures 1 and 2; Figure 3 – Supplementary Table 1**). When we superimposed their isolated transport domains on the intracellular regions (HP1, TM8b, and TM7a), “below” the substrate-binding site, they aligned well (**Figure 3a**). In contrast, we observed displacements of helices in the extracellular halves “above” the substrate-binding site, most noticeable in HP2, TM8a, and TM7b (**Figure 3a; Figure 3 – Movie 1**). These observations suggest heterogeneity in packing of the extracellular half of the transport domain immediately after substrate binding. Similar superpositions of transport domains from the previously reported structures of substrate-, inhibitor-, and Na^+^-only bound Glt_Ph_ (Alleva et al., 2020; Boudker et al., 2007; Yernool, Boudker, Jin, & Gouaux, 2004) also picture differences in positions of TM7b and TM8a in addition to the expected changes in gating HP2 (**Figure 3b**). These observations further support conformational lability of these helices and suggest that ligand binding entails the restructuring of the entire region.

**Figure 3.**
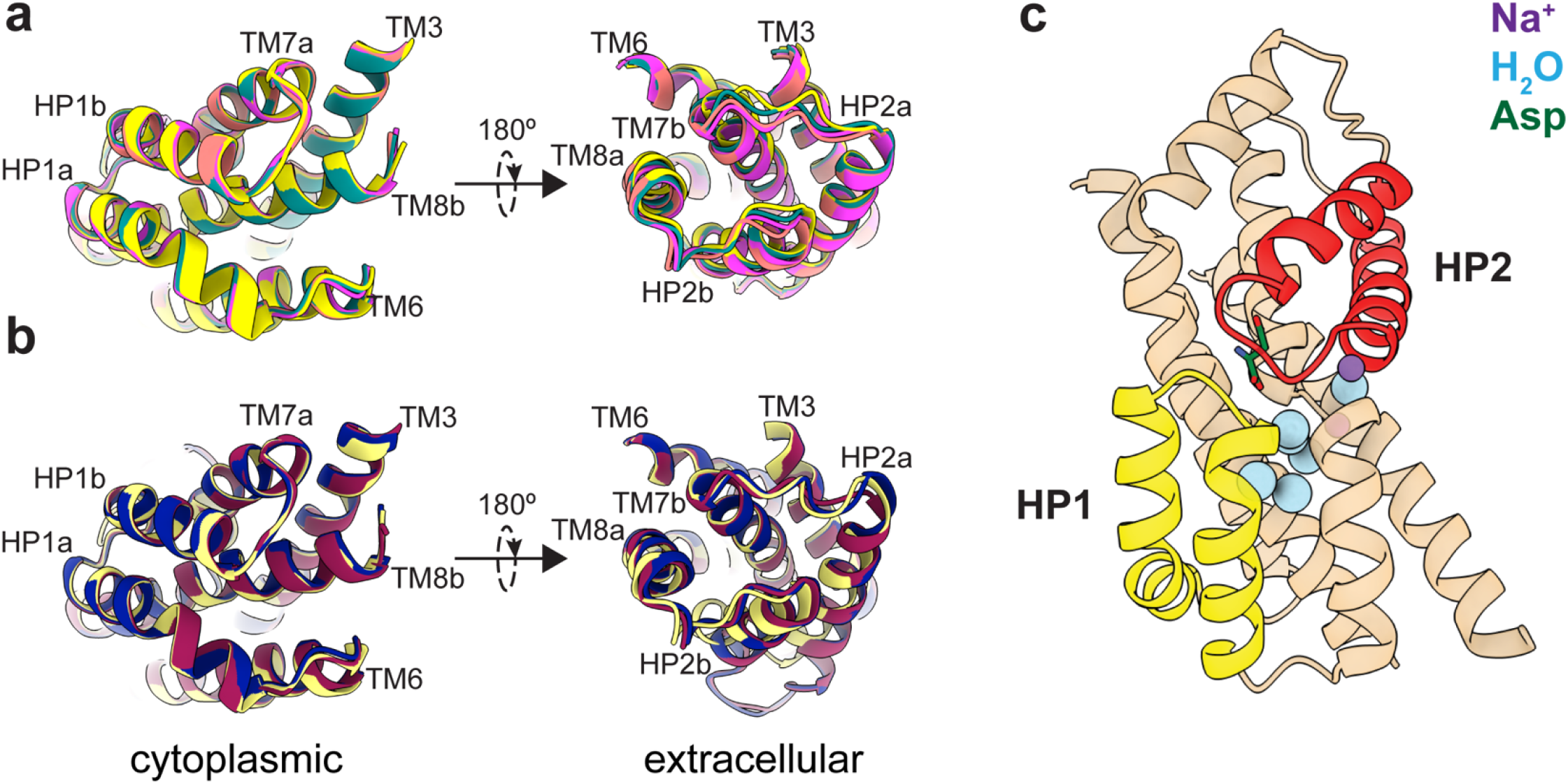
Mobility of transport domains helices. (a) Superimposition of transport domains from Dataset A. A1 is salmon, A3 teal, A6 yellow, and A7 magenta. (b) Superimposition of crystal structures of substrate-bound (2NWX; light yellow), TBOA-bound (2NWW; dark blue), and Na^+^-bound (7AHK; maroon) Glt_Ph_. The domains were superimposed on HP1 and TM7a (residues 258-309). The views are from the intracellular (left) and extracellular (right) sides of the transport domain. (c) Cartoon representation of the transport domain resolved in the equilibrated Dataset B after refinement in C3. Resolved waters are shown as blue spheres. Sodium ions are purple spheres, and the substrate is green, HP1 yellow, and HP2 red.

### High-resolution equilibrated structures of P-Glt_Ph_

ITC experiments suggest that the low-affinity OFS2 should be transient in the presence of substrate, and the high-affinity OFS1 should predominate at equilibrium (**Figure 1i**). To visualize OFS1, we imaged P-Glt_Ph_ in equilibrium conditions, purified in the presence of 250mM NaNO_3_ and 1mM L-Asp (Dataset B). We refined the maps to 2.2 Å after imposing C3 symmetry (**Figure 3 – Supplementary Figure 3**). The increased resolution compared to Dataset A may reflect reduced structural heterogeneity but can also be due to different microscopes, imaging parameters, or artifacts of grid freezing. The D405N mutation abolishes Na^+^ binding at Na1 (Boudker et al., 2007; Riederer, Moenne-Loccoz, & Valiyaveetil, 2021); in its place, we observe an excess density suggesting that a water molecule replaces the ion (**Figure 3 – Supplementary Figure 4a**). The S279E side chain points into the extracellular milieu, away from the transport domain (**Figure 3 – Supplementary Figure 4b**).

We found that the equilibrated transport domain structure is nearly identical to Class A3 (with overall r.m.s.d. of 0.30 Å). Thus, we propose that this conformation corresponds to the higher-affinity OFS1 sub-state. The ensemble of other conformations observed in Dataset A (A1, A6, A7), showing different helical packing of the transport domain, together make up the low-affinity OFS2 sub-state. This hypothesis is consistent with the observation that mutations in HP2, disrupting the hairpin packing with TM8a and TM7b helices and increasing local dynamics, V366A and A345V, decrease substrate affinity (Huysmans et al., 2021).

Interestingly, we observed several water densities within the transport domain of the equilibrated structure contributing to the hydrogen bond network between substrate- and ion-coordinating residues. All six resolved buried water molecules are below the substrate-binding site. In contrast, the extracellular half of the domain, corresponding to the labile helices in Dataset A, appears “dry” (**Figure 3c**). We speculate that the extensive hydrogen bond network in the cytoplasmic half of the transport domain ensures its rigidity. In contrast, the extracellular half, less constrained by polar interactions, can sample multiple conformations with altered packing. Notably, chaotropic salts and elevated temperature favor OFS2 consistent with less well-packed, more dynamic, water-accessible structures.

We also looked for structural heterogeneity in the Dataset B. 3D Variability (3DVA) analysis on a single protomer using the symmetry-expanded particle stack (Punjani & Fleet, 2021) revealed small movements of the transport domain (**Figure 4 – Movies 1 and 2**). Focused 3D classification of ~1.6 million symmetry-expanded particles showed only protomers in OFS and yielded structural classes corresponding to the density variations seen in 3DVA analysis. We refined these classes to 2.36-2.65 Å resolutions (**Figure 4 – Supplementary Figures 1 and 2, Figure 4 – Supplementary Table 1**). Superimpositions of the refined trimers on trimerization regions (residues 150-195) showed three subtly different tilts of the classified protomers, consisting of movements of the transport domain and the peripheral parts of the scaffold (OFS_out_, OFS_mid_, and OF_Sin_) (**Figure 4 – Movie 3**). The largest tilt difference of 2.1° is between OFS_out_ and OF_Sin_ transport domains. The tilt differences for OFS_out_/OFS_mid_ and OFS_mid_/OFS_in_ were ~1.1° each **(Figure 4 – Supplementary Figure 3**). The adjacent protomers are unaffected, suggesting that the movements occur independently in individual protomers (**Figure 4 – Movie 3; Figure 4 – Supplementary Figure 4**). Notably, we observed no rearrangements of the extracellular helices regardless of the tilts, and all tilt states most closely resembled Class A3 in Dataset A, consistent with the high-affinity OFS1 predominating at equilibrium (**Figure 4 – Supplementary Figure 5; Figure 4 – Supplementary Table 2**). The mechanistic basis of the tilts and their role in transport remain unclear. Similar transport domain tilts might also be present in Dataset A; however, the moderate resolution of the maps prevents their visualization.

Analysis of the substrate-binding sites of Dataset B classes revealed that Asp-390 sampled multiple rotameric states. The highly conserved Asp-390 in TM8 does not coordinate the substrate but is critical for the high-affinity binding – D390A mutant has a 1000-fold lower affinity (Riederer & Valiyaveetil, 2019). Arg-397 in TM8 is the principal substrate-coordinating residue. Its guanidinium group forms hydrogen bonds with the L-Asp sidechain carboxylate and cation-π interactions with Tyr-317 in TM7. We find that Asp-390 can be in “down” or “up” rotamers, hydrogen-bonding to Arg-397 or Tyr-317, respectively (**Fig. 5a-b**). Classifications also revealed a “middle” rotamer, perhaps representing an average of the two rotamers or a unique state (**Figure 5c**). Different Asp-390 rotamers do not result in an observable change of Arg-397 conformation but might alter the local electrostatics. Furthermore, tyrosine hydrogen-bonding through the OH group potentiates cation-π interactions compared to phenylalanine (Gallivan & Dougherty, 1999), and Y317F mutation leads to a 10-fold loss of L-Asp affinity in Glt_Ph_ (Riederer & Valiyaveetil, 2019). Thus, the “up” and “down” rotamers might alter substrate affinity. After additional rounds of sorting, we found that only subpopulations of OFS_out_, comprising ~10% of all particles, featured “up” or “middle” rotamers (**Figure 5c; Figure 5 – Supplementary Table 1; Figure 4 – Supplementary Figure 1**). Thus, the preference of Asp-390 to hydrogen-bond with Arg-397 or Tyr-317 might be allosterically coupled to the position of the transport domain. Whether these structural heterogeneities contribute to the elevator dynamic or substrate binding heterogeneities is yet unclear.

**Figure 5:**
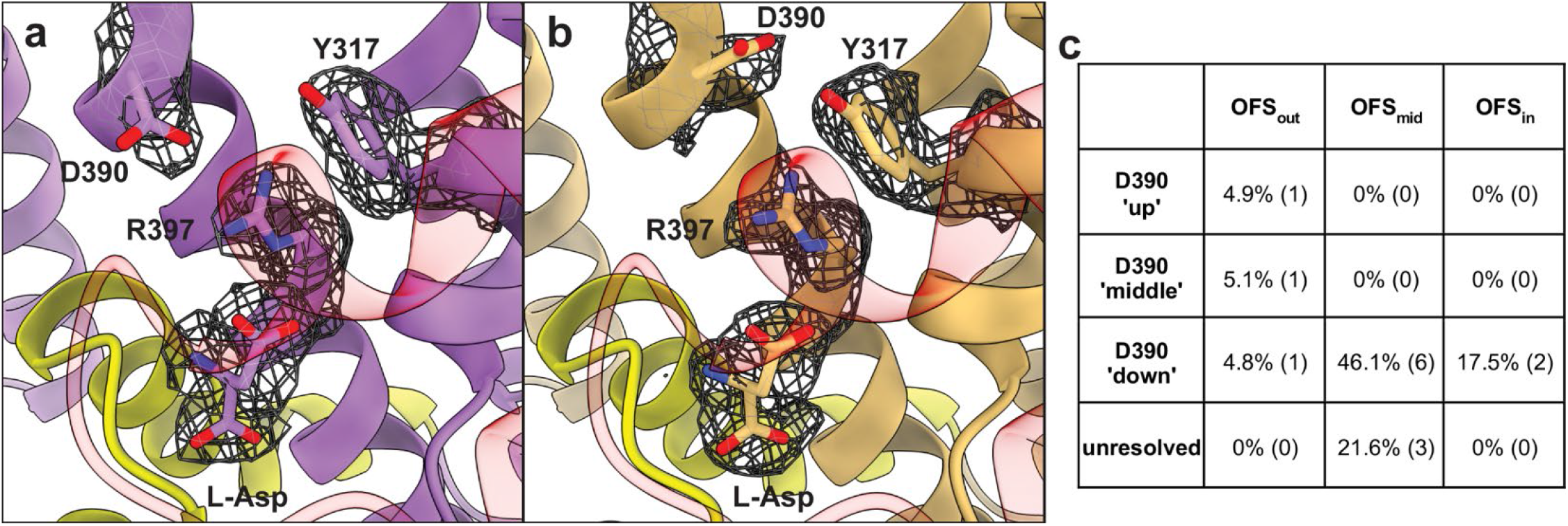
Multiple rotameric states of D390. Cartoon representations of OFS_out_ classes with “down” (a) and “up” (b) D390 rotamers. The mesh objects are density maps contoured at 4 σ; only densities within 1.5 Å of labeled residues are displayed for clarity. HP1 is in yellow, and HP2 in red. (c) Percentage of particles (total = 1,623,456) that classified into certain D390 rotamers. Numbers in parentheses represent the number of 3D classes, further detailed in Figure 4 – Supplementary Figure 1 and Figure 5 – Supplementary Figure 1.

## DISCUSSION

Serendipitously, we found that P-Glt_Ph_ mutant has two slowly-exchanging outward-facing conformational sub-states, OFS1 and 2, binding aspartate with different affinities and enthalpies (**Figure 1**). smFRET and cryo-EM showed that P-Glt_Ph_ is predominantly outward-facing (**Figure 1a-b**), possibly due to S279E mutation at the tip of HP1 (**Figure 3 – Supplementary Figure 4b**). Thus, P-Glt_Ph_ is an excellent model to consider the mechanism of heterogeneous binding in OFS. WT Glt_Ph_ is less suitable because its conformational ensemble includes iOFS and IFS (Akyuz et al., 2013; Reyes et al., 2013; X. Wang & Boudker, 2020). Nevertheless, WT binding isotherms also suggest multiple binding conformations modulated by salts and temperature (**Figure 2**). Interestingly, a recent saturation transfer difference NMR study reported an unusually low Hill coefficient of 0.69 for aspartate binding to liposome-reconstituted Glt_Ph_ (Hall et al., 2020). A Hill coefficient below 1 may reflect negative cooperativity or result from multiple binding states with distinct affinities (Cattoni, Chara, Kaufman, & Gonzalez Flecha, 2015; Sevlever, Di Bella, & Ventura, 2020; Z. X. Wang & Pan, 1996). Distinguishing these possibilities requires a kinetic approach (Cattoni, Kaufman, & Gonzalez Flecha, 2009).

Recent studies showed that Glt_Ph_, originating from a hyperthermophilic archaeon, exhibits activity “modes” at ambient temperature, where transporter sub-populations function with rates differing by orders of magnitude (Ciftci et al., 2020). Switching between the modes is rare, occurring on a timescale of hundreds of seconds. These modes were attributed to sub-populations with different elevator dynamics and intracellular substrate-release rates (Huysmans et al., 2021; Matin et al., 2020). Here, we observed that NaNO3 modulated the populations of the binding sub-states and increased the fraction of the slow transporters (**Figure 2f**). Therefore, the substrate-binding heterogeneity might contribute to the heterogeneous uptake rates or reflect different substrate-binding properties of “fast” and “slow” sub-populations. Some gain-of-function mutations, including R276S/M395R, A345V, and V366A, both increase Glt_Ph_ elevator dynamics and reduce substrate affinity (Ciftci et al., 2021; Huysmans et al., 2021). Thus, structural perturbations can affect binding and translocation in concert, suggesting that the two processes share some of the determinants. Therefore, insights into the structural bases of heterogeneous binding might also illuminate the origins of heterogeneous transport kinetics.

We thoroughly examined multiple cryo-EM datasets to understand the structural basis of substrate-binding heterogeneity. The structural heterogeneity most consistent with the binding data is the altered packing in the extracellular half of the transport domain (**Figure 3a-b**). It is observed exclusively in Dataset A, collected immediately after substrate binding to P-Glt_Ph_, and not the equilibrated transporter (Dataset B). Remarkably, one of the classes in Dataset A showed the transport domain structure identical to Dataset B, which we interpret as the high-affinity OFS1. We think the other classes in Dataset A, showing movements of helices in the extracellular half, collectively represent the low-affinity OFS2. We further suggest that the relaxation from OFS2 to OFS1 is slow because it entails repacking a large transport domain region and perhaps, even unbinding and rebinding of aspartate or sodium. Consistently, stopped-flow experiments suggested that substrate and Na^+^ binding to the transporter involves a kinetically slow step to achieve a tightly bound state, likely following sodium binding (Ewers, Becher, Machtens, Weyand, & Fahlke, 2013; Hanelt et al., 2015). Because HP2 closure upon substrate binding is rapid (Riederer & Valiyaveetil, 2019), helical repacking might be the slow step.

Our structural analysis suggests that there is a structural specialization within the transport domain of glutamate transporters. A network of buried waters and polar residues within the cytoplasmic half of the domain (TM8b, TM7a, HP1) might ensure exquisitely specific, evolutionarily conserved structure responsible for sodium selectivity and allosteric coupling between ion and substrate binding (**Figure 3c**). In contrast, the extracellular half (TN7b, HP2, TM8a) is more hydrophobic and contains no resolved waters. Fewer constraining polar interactions may permit variable helical packing in this region, setting the dynamic properties – substrate affinities and elevator dynamics – of Glt_Ph_ sub-states and, perhaps, different glutamate transporter homologues. Consistently, extensive mutagenesis in HP1 and adjacent helices did not identify gain-of-function mutants analogous to those in HP2 (Huysmans et al., 2021).

The structural differences between the high- and low-affinity states are small and require analysis of high-resolution cryo-EM datasets. Similarly, modal gating in KcsA was attributed to subtle sidechain rearrangements (Chakrapani et al., 2011). Functional heterogeneity has been observed in ion channels, transporters, and enzymes. For example, nicotinic acetylcholine receptors feature opening bursts interspersed with short or long closed periods (Colquhoun & Sakmann, 1985), and ionotropic glutamate receptors show complex kinetics with multiple gating modes (Popescu, 2012). P-type ATPase also displayed periods of rapid transport interspersed with prolonged pauses (Veshaguri et al., 2016). As in Glt_Ph_, these distinct modes may be due to concerted subtle restructuring of protein regions occurring on long timescales.

## MATERIALS AND METHODS

### DNA manipulations and protein preparation

Mutations were introduced to the previously described Glt_Ph_ CAT7 construct (Yernool et al., 2004) using PfuUltra II, and sequences were verified using Sanger sequencing (Macrogen USA). Proteins were expressed as C-terminal (His)_8_ fusions, separated by a thrombin cleavage site. Proteins were purified as previously described (Yernool et al., 2004). Briefly, crude membranes of DH10B *E. coli* cells overexpressing Glt_Ph_ were solubilized for 2 hours in 20mM HEPES/NaOH pH 7.4, 200mM NaCl, 5mM L-Asp, and 40mM n-dodecyl-ß-d-maltopyranoside (DDM, Anatrace). After solubilization, the DDM was diluted to ~8-10 mM, and after a high-speed ultracentrifugation step (40,000 rpm, Ti45 rotor), the supernatant was applied to pre-equilibrated Ni-NTA affinity resin (Qiagen) for 1 hour. The resin was washed with 7 column volumes of 20mM HEPES/NaOH pH 7.4, 200mM NaCl, 5mM L-Asp, and 40mM imidazole. Subsequently, the resin was eluted with the same buffer with increased imidazole (250mM). The (His)_8_ tag was cleaved by thrombin digestion overnight at 4°C, and the proteins were further purified by size exclusion chromatography (SEC) in the appropriate buffer for subsequent experiments. The concentration of Glt_Ph_ protomers was determined in a UV cuvette with a 10 mm pathlength (Starna Cells, Inc.), using protein diluted 1:40, and an experimentally determined extinction coefficient of 57,400 M^−1^ cm^−1^ (Reyes et al., 2013).

### Reconstitution and uptake assays

Liposomes able to maintain proton gradients were prepared using a 3:1 mixture of 1-palmitoyl-2-oleoyl-sn-glycero-3-phosphoethanolamine and 1-palmitoyl-2-oleoyl-sn-glycero-3-phospho-(1’-rac-glycerol) (POPE/POPG). Lipids were dried on the rotary evaporator for 2 hours and under vacuum overnight. The resulting lipid film was hydrated by 10 freeze-thaw cycles at a concentration of 5 mg/mL in a 50 mM potassium phosphate buffer and 100 mM potassium acetate at pH 7. The suspensions were extruded using a Mini-Extruder (Avanti) through 400 nm membranes (Whatman) 10 times to form unilamellar liposomes, then Triton X-100 was added to liposomes at a ratio of 1:2 (w/w).

P-Glt_Ph_ for reconstitution was affinity-purified, thrombin-cleaved, and purified by SEC in 20mM HEPES/Tris pH 7.4, 200mM NaCl, 1mM L-Asp, and 7mM DM. Purified protein was added to destabilized liposomes at a ratio of 1:1000 (w:w) and incubated for 30 min at 23°C. Detergent was removed with four rounds of incubation with SM-2 beads (Bio-Rad) at 80mg beads per 1 mL of liposome suspension (2 hrs at 23°C twice, overnight at 23°C once, and 2 hrs at 23°C once). Prior to use, SM-2 beads were prewashed in methanol, rinsed thoroughly with distilled water, and equilibrated in the liposome internal buffer. After detergent removal, proteoliposomes were concentrated to 50 mg/mL by ultracentrifugation at 86000 x g for 40 min at 4°C, freeze-thawed three times, and extruded through 400 nm membranes 10 times.

Uptakes were initiated by diluting reconstituted proteoliposomes 1:100 in the appropriate reaction buffer pre-incubated at 30°C. At the indicated time points, 200μL reaction aliquots were removed and diluted in 2mL of ice-cold quenching buffer (20mM HEPES/Tris pH 7, 200mM LiCl). The quenched reaction was immediately filtered through a 0.22μm filter membrane (Millipore Sigma) and washed three times with 2mL quenching buffer. Washed membranes were inserted into scintillation vials, and the membranes were soaked overnight in 5 mL Econo-Safe Counting Cocktail. Radioactivity in liposomes was measured using an LS6500 scintillation counter (Beckman Coulter).

### Isothermal Titration Calorimetry

Substrate-free P-Glt_Ph_ and Glt_Ph_ proteins were affinity-purified, thrombin-cleaved, and purified by SEC in 20mM HEPES/KOH pH 7.4, 99mM potassium gluconate, 1mM sodium gluconate, and 1mM DDM. Proteins were immediately concentrated to > 5mg/mL and diluted 2.5-fold to a final concentration of 30-50μM. When diluted, the sample was supplemented with a final concentration of 1mM DDM, 58mM HEPES/KOH pH 7.4, and an appropriate amount of sodium salt. 350μL of protein samples were degassed, equilibrated to the temperature of the experiment, and ~300 μL was loaded into the reaction cell of a small volume NanoITC (TA Instruments, Inc.), or in the case of TFB-TBOA experiments, an Affinity Auto ITC (TA Instruments, Inc.). Titrant was prepared in a buffer matching the reaction cell, except that it contained the appropriate amount of monopotassium aspartate (RPI, Inc.) and no protein or DDM. Dilution of DDM in the reaction cell over the course of the experiment had negligible effects on the injection heats, as previously noted (Boudker & Oh, 2015). 2μL of titrant was injected every 5-6 minutes, at a constant temperature, and a stirring rate of 250 rpm (125 rpm for TFB-TBOA experiments). Injection heats measured after protein was saturated with the ligand were used to determine the dilution heats subtracted from the ligand binding heats. Data were analyzed using the NanoAnalyze software (TA Instruments, Inc.) applying either the ‘Independent’ or ‘Multiple Sites’ (referred to as ‘single-state’ and ‘two-state’ throughout the text) binding models. For two-state binding, where each state is independent and non-identical, the binding polynomial can be expressed as:

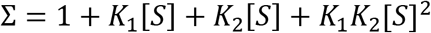

Where *K_i_*-s are the binding constants, and *[S]* is the concentration of free L-Asp. The fraction of total protein bound is given by the following:

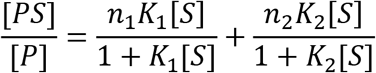

where *n_i_* is the apparent number of sites per protein molecule.

For the TFB-TBOA competition experiments, protein was first titrated with TBOA to saturation. Then, the concentration of TFB-TBOA was increased to a final concentration of 150 μM of TFB-TBOA, so that the overflow protein was also saturated. The appropriate amount of aspartate was subsequently titrated into the saturated protein to yield a binding curve.

### Cryo-electron microscopy imaging data collection

In both data sets, 3.5 μL of protein at ~4.5 mg/mL was applied to a glow-discharged QuantiFoil R 1.2/1.3, 300 mesh, gold grid (Electron Microscopy Sciences). Grids were blotted at room temperature and 100% humidity for 3 s at 0 blot force, and plunge frozen in liquid ethane using a VitroBot Mark IV (FEI). Dataset A was collected on P-Glt_Ph_, which was affinity-purified, concentrated, and buffer-exchanged into 20mM HEPES/Tris pH 7.4, 99mM K-gluconate, 1mM Na-gluconate, and 0.8mM DDM to remove the substrate. The protein was then SEC purified in 20mM HEPES/Tris pH 7.4, 250mM NaNO_3_, and 0.8mM DDM. 2.5 μL of substrate-free protein was applied to the grid, then 1μL of L-aspartate was added and mixed just prior to freezing so that the final substrate concentration was 1mM, the final protein concentration was ~4.5mg/mL, and protein was exposed to the substrate for ~5s prior to freezing (including blot time). Dataset B was collected on P-Glt_Ph_ SEC purified in 20mM HEPES/Tris pH 7.4, 250 mM NaNO_3_, 1mM L-Asp, and 0.8mM DDM. The final buffer conditions of Dataset B were identical to Dataset A.

All imaging data were collected on Titan Krios microscopes (FEI) operated at 300 kV. Dataset A was collected on a K2 Summit direct electron detector (Gatan). Automated data collection was performed in counting mode using Leginon software (Suloway et al., 2005), with a magnification of 22500x, electron exposure of 70.23 e^−^/Å^2^, 50 frames per second, a defocus range of −1.0 to −2.0 μm, and pixel size of 1.073 Å/pix. Dataset B was collected with a K3 Summit direct electron detector (Gatan). Automated data collection was performed in super-resolution counting mode using SerialEM software (Mastronarde, 2005) with a magnification of 81000x, electron exposure of 47.91 e^−^/Å^2^, 30 frames per second, a defocus range of −0.5 to −2.5 μm, and pixel size of 0.53 Å/pix.

### Image processing

The frame stacks were motion-corrected using MotionCor2 (Zheng et al., 2017), with 2x binning in the case of Dataset A, and contrast transfer function (CTF) estimation was performed using CTFFIND 4.1 (Rohou & Grigorieff, 2015). All datasets were processed using cryoSPARC 3.0 and Relion 3.0.8 simultaneously with default parameters unless otherwise stated (Punjani, Rubinstein, Fleet, & Brubaker, 2017; Su et al., 2020; Zivanov et al., 2018). Specific information on the processing of each dataset is in **Figure 2 – Supplementary Figures 2-4**. In brief, particles were non-specifically picked from micrographs using the Laplacian-of-Gaussian (LoG) picker, aiming for ~2000 picks per micrograph. These particles were extracted at a box size of 240 pixels with 4x binning. These particles were then imported to cryoSPARC, underwent one round of 2D classification to remove artifacts, and then multiple rounds of heterogeneous refinement (C1) using eight total classes, seven of which were noisy volumes (created by one iteration of *ab initio*) and one of which were an unmasked 3D model obtained from a previous processing pipeline. Once > 95% of particles converged on a single class, the particles were converted back to Relion format via PyEM (Asarnow, Palovcak, & Cheng, 2019), and re-extracted at full box size. These particles were reimported to cryoSPARC, underwent three more rounds of heterogeneous refinement, then non-uniform (NU) refinement using C3 symmetry (dynamic mask threshold, dynamic mask near, and dynamic mask far were always set to 0.8, 12, and 28, respectively) (Punjani, Zhang, & Fleet, 2020). These particles were converted back to Relion format and underwent Bayesian polishing, using parameters obtained using 5,000 random particles within the set. After three more rounds of heterogeneous refinement in cryoSPARC and one round of NU-refinement, we performed local CTF refinement (minimum fit res 7 Å). After three more rounds of heterogeneous refinement and one round of NU-refinement, we performed global CTF refinement (3 iterations, minimum fit res 7 Å; fitting trefoil, spherical aberration, and tetrafoil), three more rounds of heterogeneous refinement, and one round of NU-refinement.

Dataset B was also further processed to obtain the high-resolution structure by three additional rounds of polishing, local CTF refinement, and global CTF refinement as described above. During the polishing rounds, the box size and pixel size was rescaled as indicated in the supplement. After the second round of polishing, we classified single protomers by employing the ‘relion_particle_symmetry_expand’ function in Relion to artificially expand the particle set three times (C3) so that each protomer rotated to the same position (Scheres, 2016). The expanded particle set was subjected to 3D classification without alignment with T=400 and 10 classes, using the refined C3 structure as a reference map. The exceptionally high T value was chosen to separate out subtle structural changes, and lower T values were also tested during processing. The local mask was created using a 20 Å map of the transport domain of Chain A of PDB model 2NWX, with an initial binarization threshold of 0.01, extended by 3 pixels, and a soft-edge of 10 pixels. Of these, particle stacks from subsets of interest were separately used in cryoSPARC’s local refinement (C1), using the mask and map obtained from the most recent NU-refinement. Single protomers in Dataset A were classified similarly, except we performed symmetry expansion after the first round of polishing due to the limited resolution of the dataset. After processing, the resulting half-maps were used as inputs for density modification implemented in PHENIX 1.19.1-4122 (Adams et al., 2010; Terwilliger, Ludtke, Read, Adams, & Afonine, 2020), using a mask created from the NU-refinement job (threshold 0.1, dilation radius 15, soft padding width 5). All density maps were displayed using ChimeraX (Pettersen et al., 2021).

### Model building and refinement

For atomic model building, the crystal structure of WT Glt_Ph_ in the OFS (2NWX) was docked into the densities using UCSF Chimera (Pettersen et al., 2004). The model was first real-space refined in PHENIX (Adams et al., 2010). Then, chain A was adjusted manually, and ions were added in COOT (Emsley & Cowtan, 2004). Waters were initially added using phenix.douse, and subsequently manually inspected and adjusted. The resulting model underwent additional rounds of real-space refinement and validated using MolProbity (Chen et al., 2010). All structural models were displayed using ChimeraX (Pettersen et al., 2021). Per-residue Cα RMSDs were generated with Chimera (Meng, Pettersen, Couch, Huang, & Ferrin, 2006). To cross-validate models, refined models (FSC_sum_) were randomly displaced an average of 0.3 Å using phenix.pdbtools. The displaced model was real-space refined against half-map 1 obtained through density modification to obtain FSC_work_. The resulting model was validated against half-map 2 to obtain FSC_free_.

### Single-molecule dynamics assay

Thrombin-cleaved P-Glt_Ph_, containing C321A and N378C mutations, was labeled and reconstituted as described previously (Akyuz et al., 2013). Protein was SEC purified in 20 mM HEPES/Tris, 200 mM NaCl, 1 mM L-Asp, and 1 mM DDM. Purified protein was labeled using maleimide-activated LD555P-MAL and LD655-MAL dyes and biotin-PEG11 at a molar ratio of 4:5:10:2.5. Excess dyes were removed on a PDMiniTrap Sephadex G-25 desalting column (GE Healthcare).

All experiments were performed on a previously described home-built prism-based TIRF microscope constructed around a Nikon Eclipse Ti inverted microscope body (Juette et al., 2016). Microfluidic imaging chambers were passivated with biotin-PEG, as previously described (Akyuz et al., 2015). After passivation, the microfluidic channel was incubated with 0.8 μM streptavidin (Invitrogen) in T50 buffer (50 mM NaCl, 10 mM Tris, pH 7.5) for 7 min, then thoroughly rinsed with T50 buffer. Detergent-solubilized protein was immobilized by slowly flowing over the channel, and excess protein was removed by washing with 1 mL 25 mM HEPES/Tris pH 7.4, 200 mM KCl, 1 mM DDM.

Buffers were supplemented with an oxygen-scavenging system composed of 2 mM protocatechuic acid (PCA) and 50 nM protocatechuate-3,4-dioxygenase (PCD), as described previously (Aitken, Marshall, & Puglisi, 2008). The smFRET movies were recorded with 100 ms integration time using 80-100 mW laser power. All conditions tested were in the presence of 25 mM HEPES/Tris, 500 mM sodium salt, and 1 mM DDM, either in the presence or absence of 1 mM L-Asp. Slides were washed with 25 mM HEPES/Tris, 200 mM KCl, 1 mM DDM between experiments. Traces were analyzed using the Spartan software (Juette et al., 2016). Trajectories were corrected for spectral crosstalk and preprocessed automatically to exclude trajectories that lasted 15 or fewer frames and had a signal-to-noise ratio of 8 or lower. Traces with multiple photobleaching events (indicative of multiple sensors in a protein) or inconsistent total fluorescence intensity were also discarded.

### Single-molecule transport assay and analysis

Thrombin-cleaved Glt_Ph_ (C321A/N378C) was SEC purified in 20 mM HEPES/Tris pH 7.4, 200 mM NaCl, and 0.1 mM L-Asp. Protein was labeled with maleimide-activated biotin-PEG_11_ (EZ-Link, ThermoFisher Scientific) in the presence of *N*-ethylmaleimide (NEM) at a molar ratio of 1:2:4 as previously described (Ciftci et al., 2020). Liposomes were prepared from a 3:1 (w:w) mixture of *E. coli* polar lipid extract (Avanti Polar Lipids, Inc.) and egg phosphatidylcholine in SM-KCl buffer (50 mM HEPES/Tris pH 7.4, 200 mM KCl). Liposomes were extruded through 400-nm filters (Whatman Nucleopore) using a syringe extruder (Avanti), and destabilized by the addition of Triton X-100 at 1:2 (w:w) detergent-to-lipid ratio. Labeled Glt_Ph_ was added to the liposome suspension at 1:1000 (w:w) protein-to-lipid ratio at room temperature for 30 min. Detergent was removed by six rounds incubation of Bio-Beads (two rounds at 23°C for 2 hours each, one round at 4°C overnight, three rounds at 4°C for 2 hours each). The excess substrate was removed by three rounds of: centrifugation for 1 hour at 49,192 x g at 4°C, removal of the supernatant, the addition of 1mL fresh SM buffer, and three freeze/thaw cycles. Liposomes were concentrated to 50 mg/mL, and ccPEB1a-Y198F labeled with activated LD555P-MAL and LD655-MAL dyes as described (Ciftci et al., 2020) was added at a final concentration of 0.6 μM and encapsulated by two freeze-thaw cycles. To remove unencapsulated ccPEB1a-Y198F, 1 mL of SM-KCl buffer was added, and liposomes were centrifuged for 1 hour at 49.192 x g at 4°C. Supernatants were discarded, liposomes were suspended at 50 mg/mL, and extruded 12 times through 100-nm filters.

Single-transporter smFRET assays were performed on the same microscope described above, and the microfluidic imaging chambers were prepared in the same way. After coating with streptavidin, the channel was rinsed thoroughly with SM-K(X) buffer (50 mM HEPES/Tris pH 7.4 containing 200 mM potassium salt buffer, where the anion (X) was changed based on the condition tested). Extruded liposomes were immobilized by slowly flowing over the channel, and excess liposomes were removed by washing with 1 mL SM-K(X) buffer.

Buffers were supplemented with an oxygen-scavenging system as above, and the smFRET movies were recorded with 400 ms integration time using 20-40 mW laser power. To confirm liposomes were not leaky, a movie was taken in SM-K(X) buffer containing 1 μM L-Asp and 1 μM valinomycin. No L-Asp uptake was observed under these conditions lacking Na^+^ gradient. After completion of this movie, another movie was initiated to record transport events. At approximately 3 seconds into this movie, SM-Na(X) buffer containing 1 μM L-Asp and 1 μM valinomycin was perfused into the channel.

Traces were analyzed using the Spartan software (Juette et al., 2016). Trajectories were corrected for spectral crosstalk and preprocessed automatically to exclude trajectories that lasted 15 or fewer frames, had a signal-to-noise ratio of 8 or lower, and had an initial FRET efficiency less than 0.4 or greater than 0.7. Traces with multiple photobleaching events (indicative of multiple sensors in a liposome) or inconsistent total fluorescence intensity were also discarded. Remaining traces were sorted by either the presence or absence of observable transport events (responding and non-responding traces, respectively), determined by an increase in FRET efficiency from ~0.55-0.6 to ~0.75-0.8.

Responding traces from each movie were plotted as time-dependent mean FRET efficiency. Buffer replacement time was determined as described (Ciftci et al., 2020). Within each dataset, the data was normalized so that 0% is the first time point, and 100% is the first time point + 0.2 (the change in FRET efficiency upon saturation of ccPEB1a-Y198F). The resulting normalized data were multiplied by the fraction of the responding traces relative to the total traces. Data from three independent reconstitutions were merged and fitted to a tri-exponential function in GraphPad Prism 8.4.2, where Y0 = 0, Plateau = 1, percentages were set between 0 and 100, and all rate constants were set to be shared between all data sets and greater than zero.

## Supporting information

Dataset B (high resolution structure)

Dataset B (structural classes)

## ACKNOWLEDGEMENTS

We thank Drs. Eva Fortea, Maria Falzone, Philipp Schmidpeter, Biao Qiu, and Xiaoyu Wang for helpful discussions on cryo-EM data processing. We especially thank Navid Paknejad for in-depth assistance with optimizing cryo-EM processing workflows. Also, we thank Bryce Delgado for preliminary ITC experiments, Will Eng for protein expression, and Vishnu Ghani for preparation of the PEB1a protein. We thank Dr. Scott Blanchard and members of the Blanchard lab for support and resources for smFRET experiments. Finally, we thank Drs. Erika Riederer and Francis Valiyaveetil for helpful discussions and exploratory experiments. The work was supported by NIH F32 NS102325 (to KDR), AHA 19PRE34380215 (to DC), and NIH R01NS064357 and R37NS085318 (to OB). Dataset A was collected with assistance from Carolina Hernandez at the Simons Electron Microscopy Center and National Resource for Automated Molecular Microscopy located at the New York Structural Biology Center, supported by grants from the Simons Foundation (349247), NYSTAR, and the NIH National Institute of General Medical Sciences (GM103310) with additional support from Agouron Institute (F00316) and NIH S10 OD019994-01. Dataset B was collected at the UMass cryo-EM facility with help from Dr. Kangkang Song and Dr. Chen Xu.

## AUTHOR CONTRIBUTIONS

K.D.R. and O.B. designed the experiments, analyzed the data, refined the molecular models, and wrote the manuscript with input from all authors. K.D.R. and A.S. performed the cloning and radioactive transport assays. K.D.R. and D.C. performed the smFRET dynamics and transport assays. K.D.R. performed the ITC and cryo-EM sample preparation and processing.

## SUPPLEMENTARY INFORMATION

**Figure 1 - Supplementary Figure 1.**
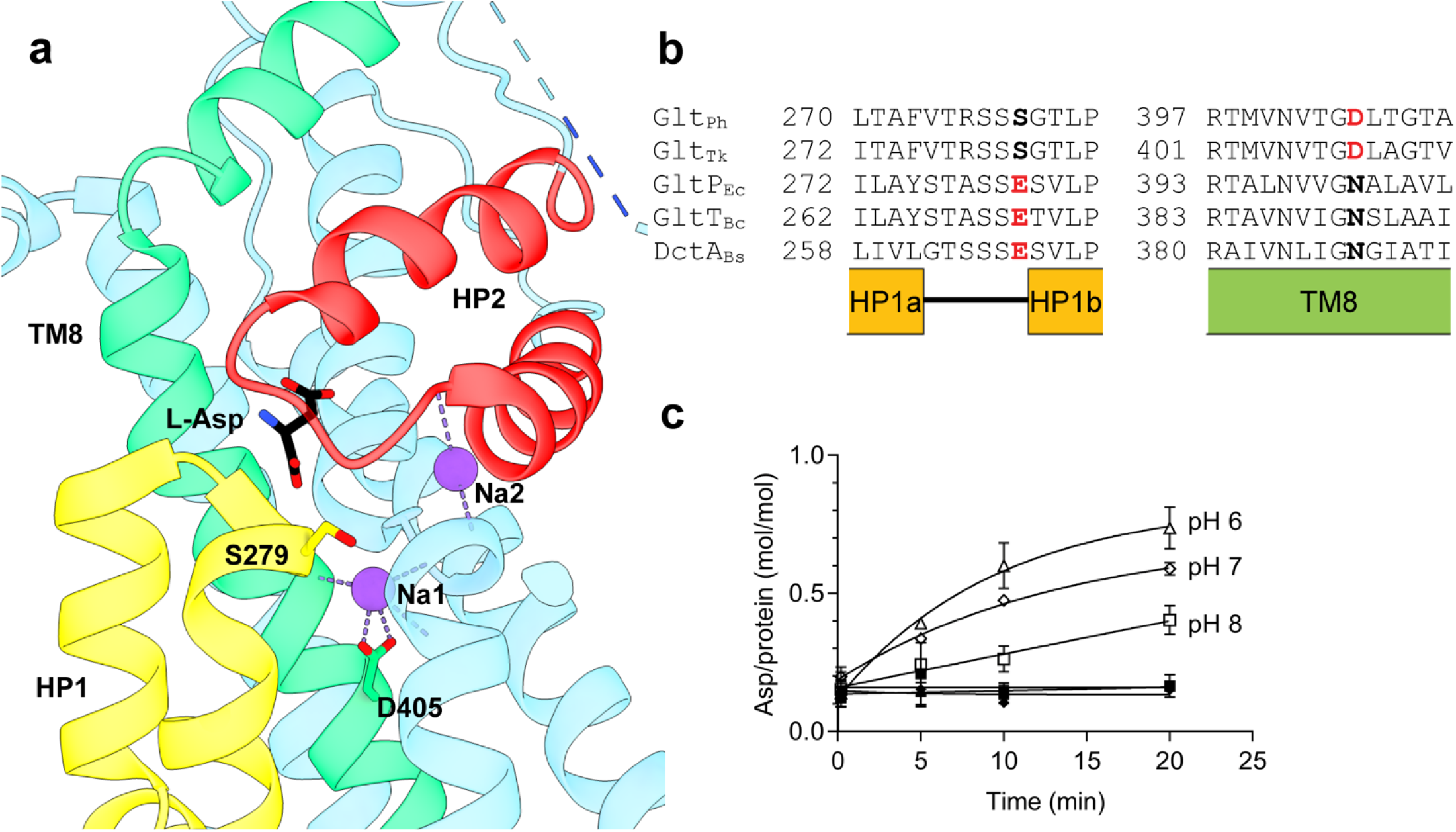
P-Glt_Ph_ (S279E/D405N) has partial proton dependence. (a) Structural model of substrate-bound Glt_Ph_ (PDB ID 2NWX). (b) Sequence alignment of Na^+^-coupled (Glt_Ph_, Glt_Tk_) and H^+^-coupled (GltP_Ec_, GltT_Bc_, DctA_Bs_) transporters; structural elements are indicated below the alignment. Residues mutated in P-Glt_Ph_ are in bold. (c) pH-dependent aspartate uptake of P-Glt_Ph_. Proteoliposomes were loaded with 50 mM potassium phosphate buffer, pH 7, and 100 mM potassium acetate and diluted into the following 50 mM buffers containing 1 μM ^3^H-L-Asp: MES/NMDG pH 6 (triangles), HEPES/Tris pH 7 (diamonds), or HEPES/Tris pH 8 (squares). Buffers contained either 100 mM KCl (filled symbols) or 100 mM NaCl (empty symbols). Solid lines are shown to guide the eye, and error bars (SD) not displayed represent errors smaller than the size of the symbol.

**Figure 1 – Supplementary Table 1.** L-Asp binding to P-Glt_Ph_ (S279E/D405N) at 10°C in 500 mM NaCl. Binding parameters are from fits to the two-state model. Averaged values are means and standard deviations from three independent experiments.

| *Trial* | $n_1$ | $K_{D,1} (M)$ | $\Delta G_1$<br>(kcal<br>mol<sup>-1</sup>) | $\Delta H_1$<br>(kcal<br>mol<sup>-1</sup>) | $\Delta S_1$<br>(cal<br>mol<sup>-1</sup> K<sup>-1</sup>) | $n_2$ | $K_{D,2} (M)$ | $\Delta G_2$<br>(kcal<br>mol<sup>-1</sup>) | $\Delta H_2$<br>(kcal<br>mol<sup>-1</sup>) | $\Delta S_2$<br>(cal<br>mol<sup>-1</sup> K<sup>-1</sup>) | % $n_2$ |
| --- | --- | --- | --- | --- | --- | --- | --- | --- | --- | --- | --- |
| **1** | 0.61 | 2.3e-9 | -11.35 | -2.06 | 32.2 | 0.13 | 1.3e-7 | -8.98 | -6.00 | 10.3 | 18 |
| **2** | 0.64 | 1.0e-10 | -13.15 | -1.64 | 40.0 | 0.20 | 5.5e-9 | -10.81 | -4.73 | 21.1 | 24 |
| **3** | 0.78 | 3.6e-9 | -11.10 | -2.13 | 31.1 | 0.18 | 1.1e-7 | -9.07 | -6.35 | 9.4 | 19 |
| **avg** |  |  | **-11.86**<br>**± 0.91** | **-1.94**<br>**± 0.22** | **34.4**<br>**± 3.9** |  |  | **-9.62**<br>**± 0.84** | **-5.70**<br>**± 0.69** | **13.6**<br>**± 5.3** | **20**<br>**± 3** |

**Figure 1 – Supplementary Table 2.** L-Asp binding to P-Glt_Ph_ (S279E/D405N) at 15°C in 500 mM NaCl. Binding parameters were fitted to the two-state model. Averaged values are means and standard deviations from three independent experiments.

| *Trial* | *n*<sub>1</sub> | *K*<sub>D,1</sub> (M) | $\Delta G_1$<br>(kcal<br>mol<sup>-1</sup>) | $\Delta H_1$<br>(kcal<br>mol<sup>-1</sup>) | $\Delta S_1$<br>(cal<br>mol<sup>-1</sup> K<sup>-1</sup>) | *n*<sub>2</sub> | *K*<sub>D,2</sub> (M) | $\Delta G_2$<br>(kcal<br>mol<sup>-1</sup>) | $\Delta H_2$<br>(kcal<br>mol<sup>-1</sup>) | $\Delta S_2$<br>(cal<br>mol<sup>-1</sup> K<sup>-1</sup>) | % *n*<sub>2</sub> |
| --- | --- | --- | --- | --- | --- | --- | --- | --- | --- | --- | --- |
| 1 | 1.13 | 1.9e-9 | -11.51 | -3.05 | 29.4 | 0.39 | 3.9e-8 | -9.77 | -6.63 | 10.9 | 26 |
| 2 | 0.64 | 1.3e-9 | -11.73 | -3.31 | 29.2 | 0.22 | 6.1e-8 | -9.51 | -7.13 | 8.3 | 26 |
| 3 | 0.63 | 3.4e-9 | -11.17 | -2.11 | 31.4 | 0.29 | 1.7e-8 | -10.23 | -9.04 | 4.2 | 31 |
| avg |  |  | -11.47<br>± 0.23 | -2.82<br>± 0.52 | 30.0<br>± 1.0 |  |  | -9.84<br>± 0.30 | -7.60<br>± 1.04 | 7.8<br>± 2.8 | 28<br>± 3 |

**Figure 2 - Supplementary Figure 1.**
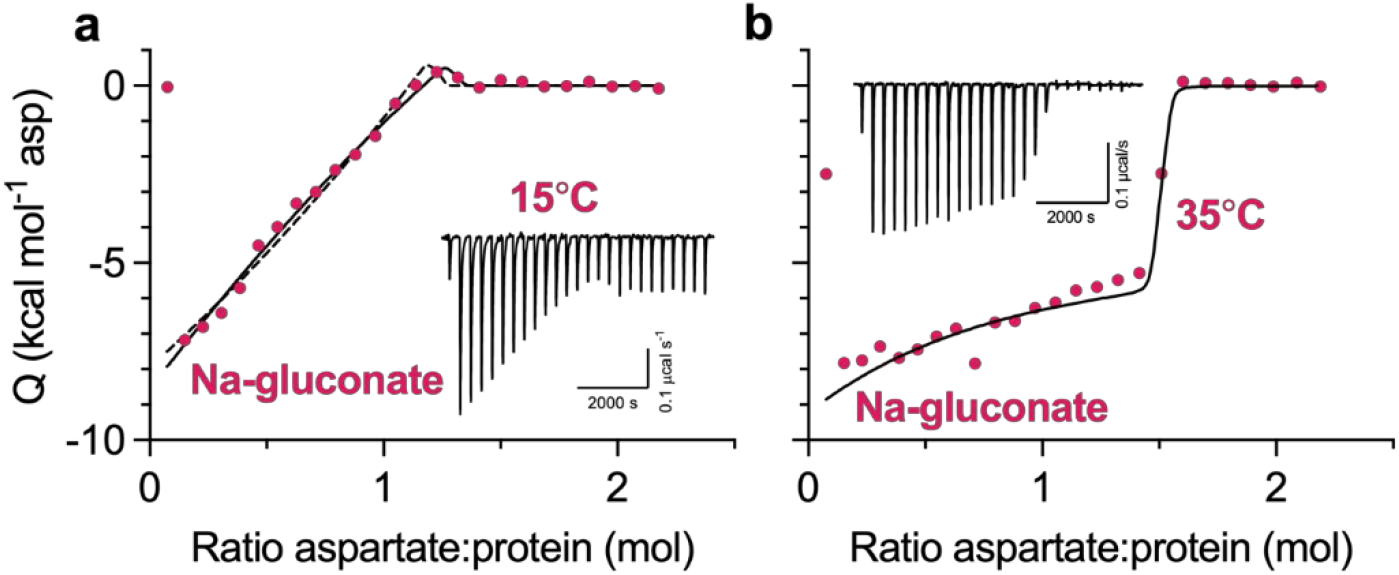
Simulated fits of wild-type binding isotherms. (a) WT Glt_Ph_, 500mM Na-gluconate, 15°C. k_D_1 and k_D_2 of Fit 1 (solid) and Fit 2 (dashed) are constrained to 0.3 nM and 0.4 nM, respectively. Fit 1 and Fit 2 have parameters of *n_1_:* 0.82 and 0.37; *ΔH_1_*: −22.7 and −50.0 kcal mol^−1^; *n_2_*: 0.36 and 0.89; *ΔH_2_*: 39.4 and 16.2 kcal mol^−1^. (b) WT Glt_Ph_, 500mM Na-gluconate, 35°C. k_D_1 and k_D_2 are constrained to 1.6 nM and 3.6 nM, respectively. Fit has parameters of *n_1_*: 0.16; *ΔH_1_*: −24.6 kcal mol^−1^; *n_2_*: 1.30; *ΔH_2_*: −4.7 kcal mol^−1^.

**Figure 2 - Supplementary Figure 2.**
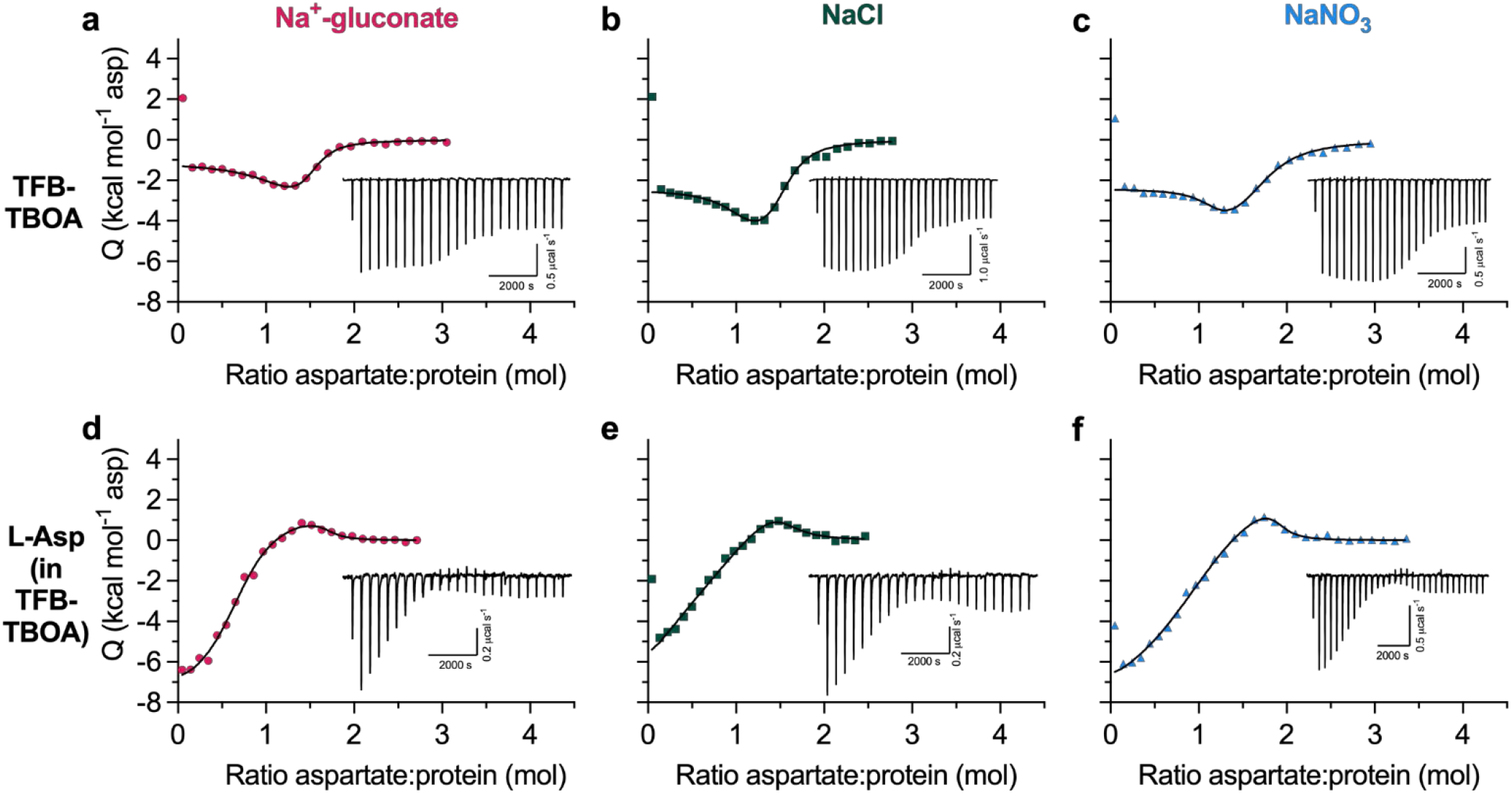
TFB-TBOA binds to two states in WT Glt_Ph_. All ITC experiments were performed at 15°C in buffers containing 500 mM Na-gluconate (red circles), NaCl (green squares), or NaNO_3_ (blue triangles). Insets show the thermal power with the corresponding scales. All data were fitted to the two-state model; however, exact binding parameters cannot be reliably determined. (a-c) TFB-TBOA binding isotherms. (d-f) Aspartate competition isotherms in the presence of saturating TFB-TBOA concentrations (Methods).

**Figure 2 - Supplementary Figure 3.**
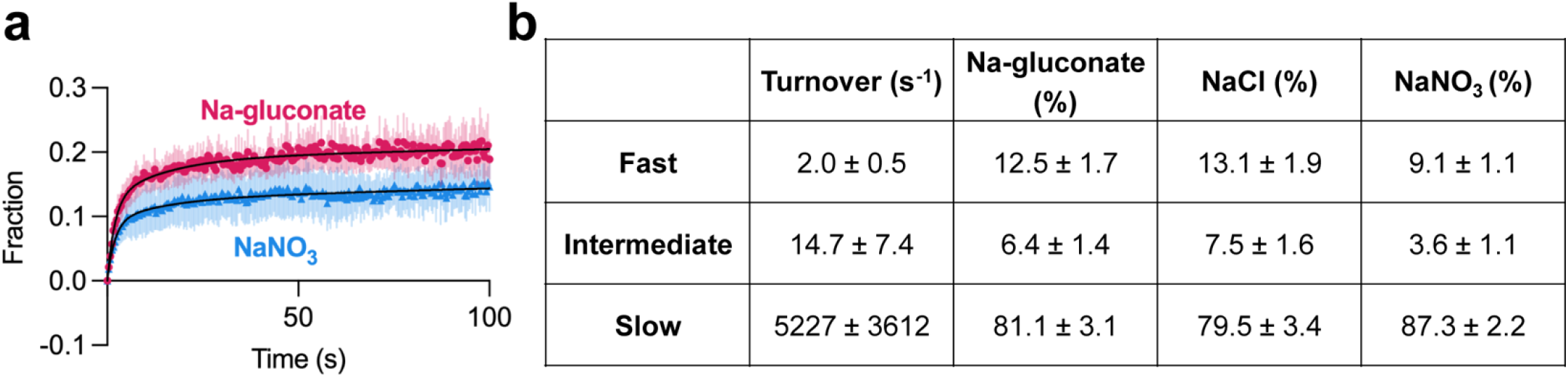
Fitted parameters of L-Asp uptake by WT Glt_Ph_ in the presence of various anions. (a) Aspartate transport of Glt_Ph_ (C321A/N378C) was measured using the single-transporter FRET-based assay in NaNO_3_ (blue) or Na-gluconate (red). Data are means and SE from three independent experiments performed as in Figure 2f. (b) Data in (a) and Figure 2f were fitted to tri-exponential functions (a three-phase association model, GraphPad Prism). Initial and final fractions of total possible uptake were set to 0 and 1, respectively. The three turnover rates (fast, intermediate, and slow), corresponding to the heterogeneous transporter populations, were constrained to be the same for all datasets (see *Methods* for a detailed description of data processing). Three independent experiments per condition were analyzed, and shown values are means and SE of the fitted parameters.

**Figure 3 - Supplementary Figure 1.**
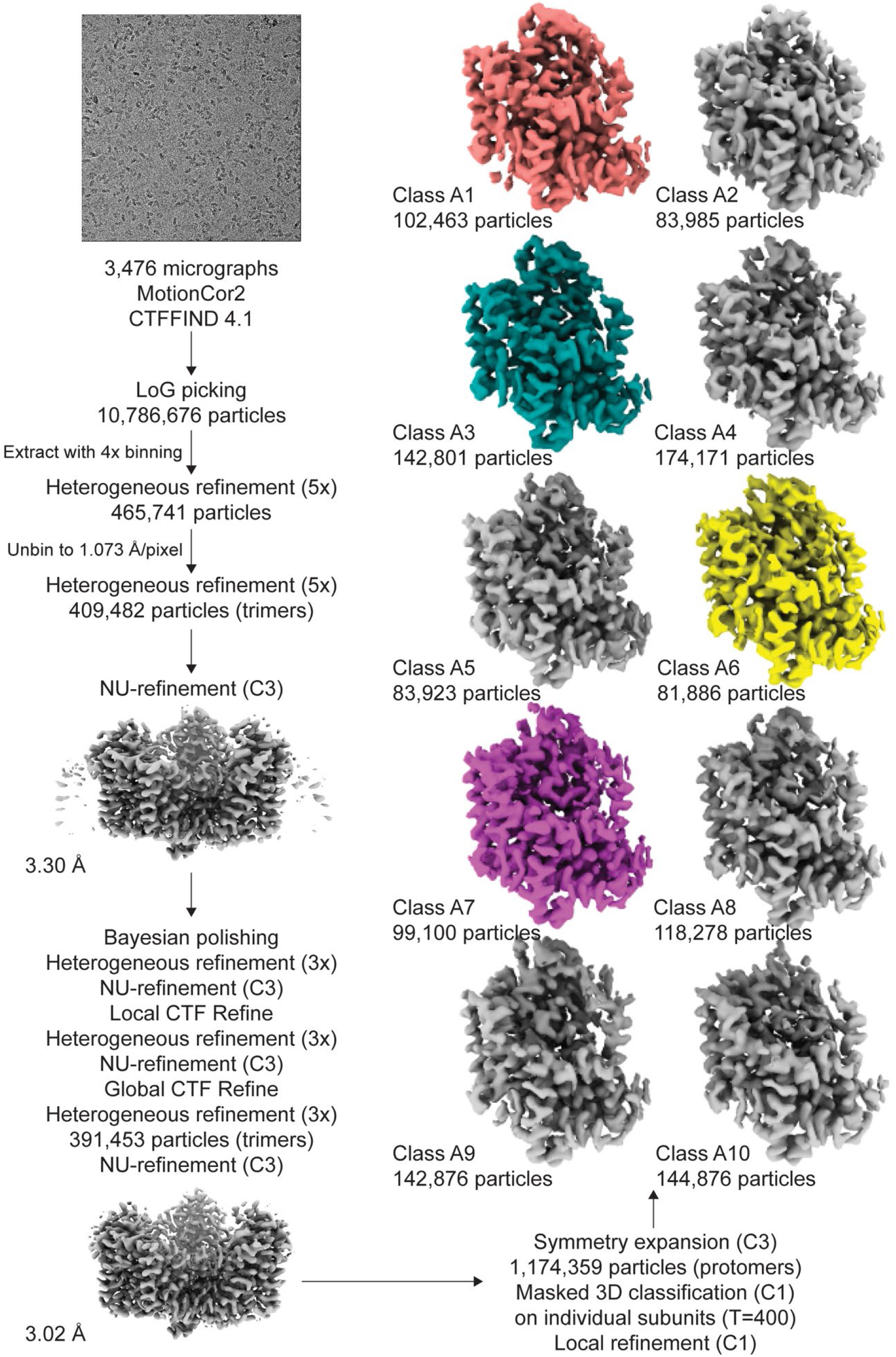
Processing flowchart for Dataset A. Protomer maps from masked classification are unsharpened, with the two other protomers removed for clarity. All maps are contoured at 8 σ.

**Figure 3 – Supplementary Table 1.** Model refinement and validation statistics for Dataset A.

|  | P-Glt<sub>Ph</sub><br>A1 | P-Glt<sub>Ph</sub><br>A3 | P-Glt<sub>Ph</sub><br>A6 | P-Glt<sub>Ph</sub><br>A7 |
| --- | --- | --- | --- | --- |
| **Cryo-EM acquisition and processing** |  |  |  |  |
| EMDataBank ID | EMD-24406 | EMD-24406 | EMD-24406 | EMD-24406 |
| Symmetry imposed | C1 | C1 | C1 | C1 |
| Initial particles | 1,174,359 | 1,174,359 | 1,174,359 | 1,174,359 |
| Class number | A1 | A3 | A6 | A7 |
| Final particles (protomers) | 102,463 | 142,081 | 81,886 | 99,100 |
| Resolution (masked FSC = 0.143, Å) | 3.42 | 3.06 | 3.65 | 3.53 |
| Density modified CC (0.5, Å) | 3.45 | 3.09 | 3.64 | 3.57 |
| **Model Refinement** |  |  |  |  |
| Model resolution (FSC = 0.50/0.143 Å) | 3.55 / 3.08 | 3.15 / 2.63 | 3.85 / 3.35 | 3.76 / 3.32 |
| **Model composition** |  |  |  |  |
| Non-hydrogen atoms | 3097 | 3094 | 3094 | 3094 |
| Protein residues | 416 | 416 | 416 | 416 |
| Ligands | 2 | 2 | 2 | 2 |
| **R.m.s. deviations** |  |  |  |  |
| Bond lengths (Å) | 0.003 | 0.002 | 0.003 | 0.003 |
| Bond angles (°) | 0.486 | 0.462 | 0.578 | 0.523 |
| **Validation** |  |  |  |  |
| MolProbity score | 1.42 | 1.21 | 1.58 | 1.78 |
| Clash score | 6.93 | 4.41 | 10.55 | 10.55 |
| Poor rotamers (%) | 0 | 0 | 0 | 0 |
| **Ramachandran plot** |  |  |  |  |
| Favored (%) | 97.82 | 98.79 | 97.82 | 96.37 |
| Allowed (%) | 2.18 | 1.21 | 2.18 | 3.63 |
| Disallowed (%) | 0 | 0 | 0 | 0 |

**Figure 3 - Supplementary Figure 2.**
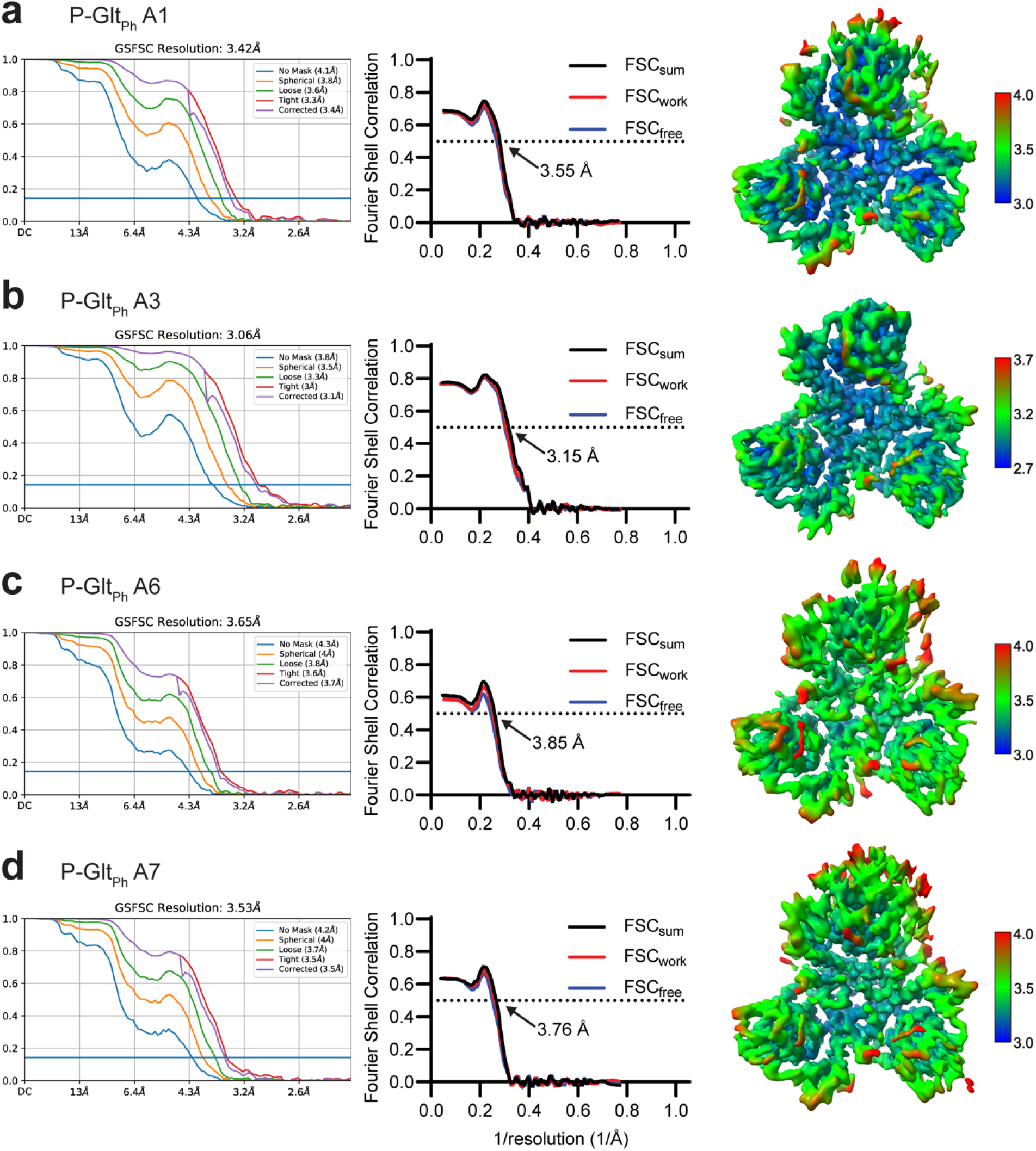
Validation statistics for models from Dataset A. From left to right, map FSC from NU-refinement in cryoSPARC, model-to-data validation in Phenix of the single protomer, and local resolution estimation of the unsharpened map. All maps are contoured at 8 σ. The top protomer is the subject of focused classification and model refinement.

**Figure 3 - Supplementary Figure 3.**
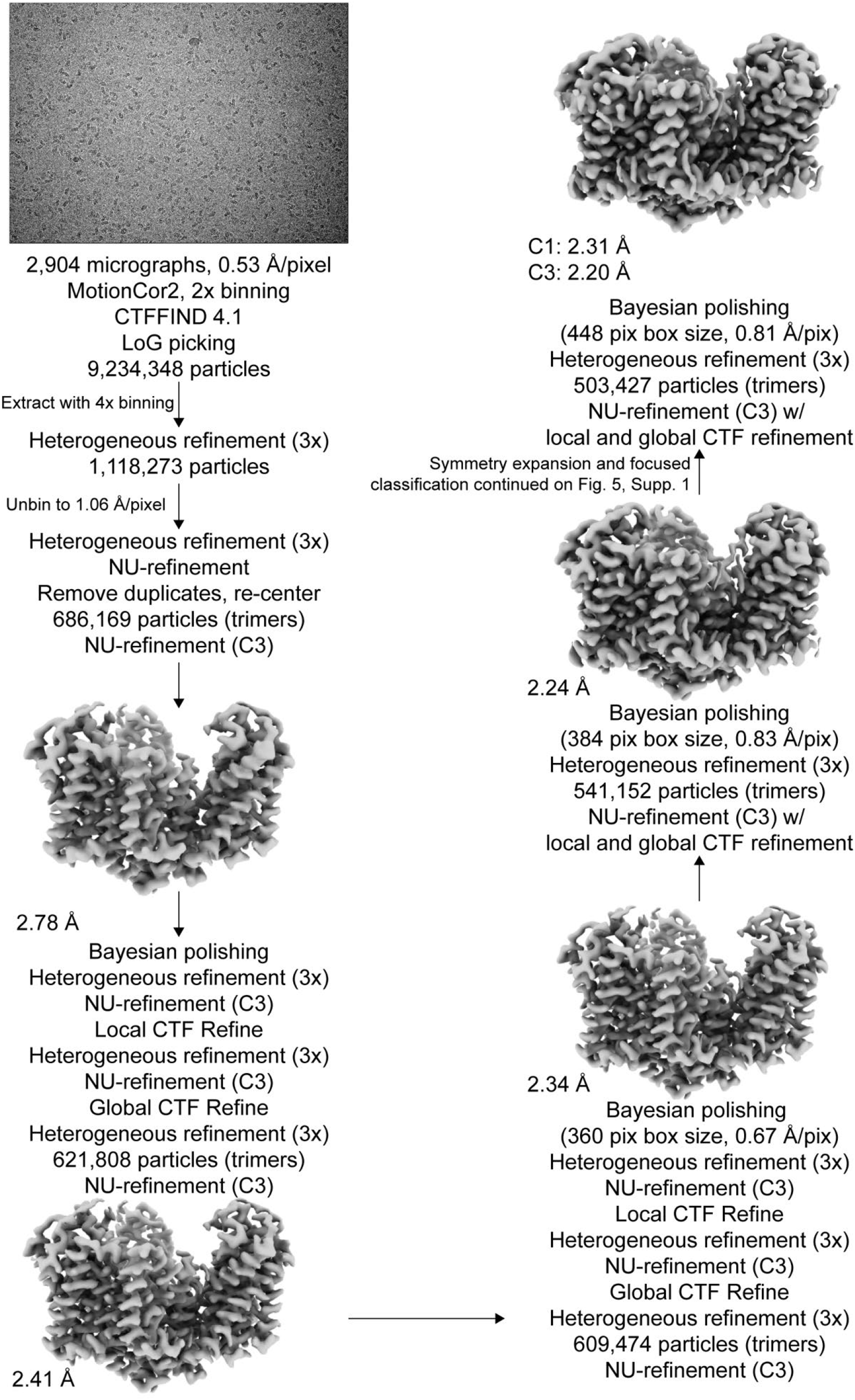
Processing flowchart for Dataset B. Unsharpened maps are contoured at σ of 8.

**Figure 3 – Supplementary Figure 4.**
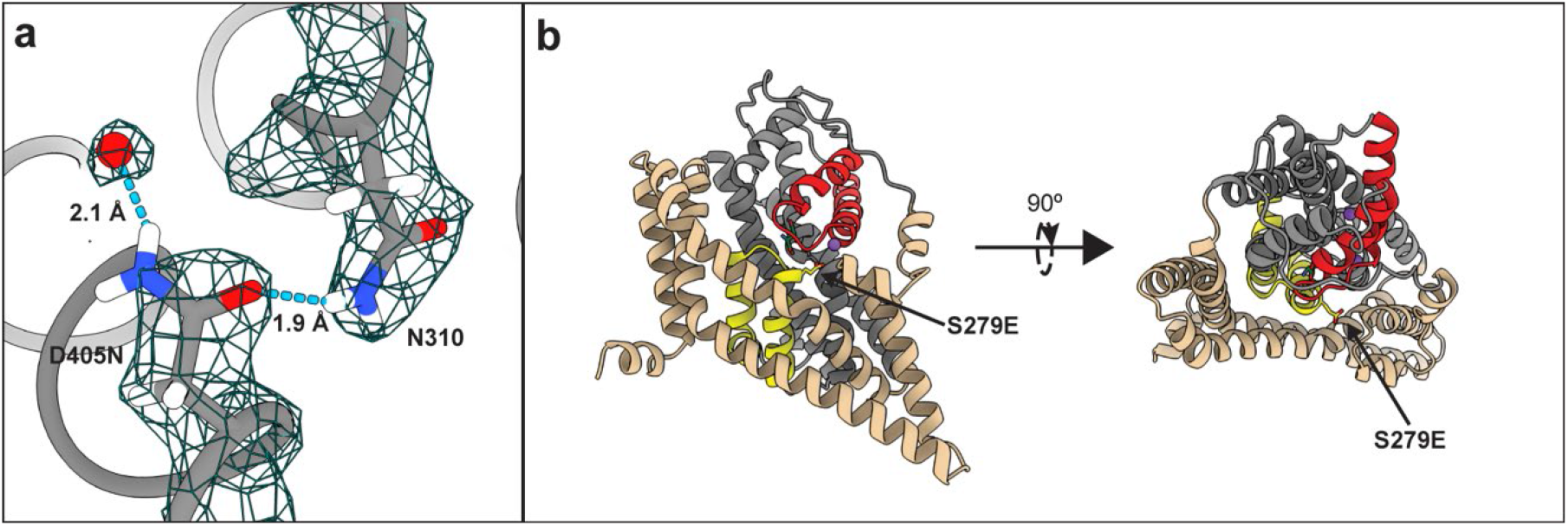
Effects of mutations on P-Glt_Ph_. (a) Close-up view of S279 mutation. The model and density map are from Dataset B refined in C3 (PDB ID 7RCP). A water molecule replacing Na1 and coordinated by D405N is emphasized as a red ball. Hydrogen bonds are displayed using ‘hbond’ in ChimeraX. (b) Cartoon representation of a protomer viewed in the membrane plane (left) and from the extracellular space (right), showing S279E points away from the transport domain.

**Figure 4 - Supplementary Figure 1.**
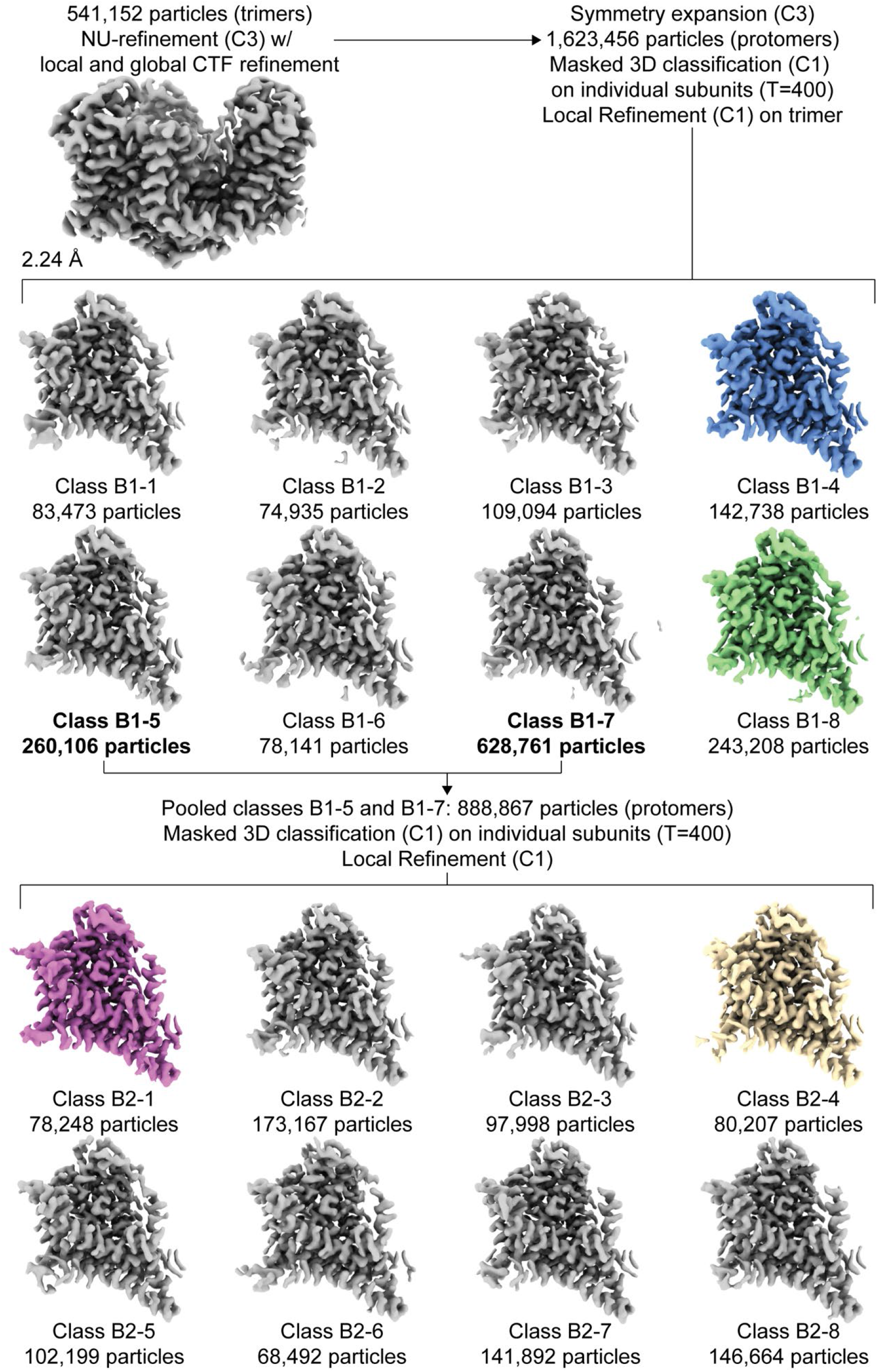
Focused classification of dataset B. Protomer maps from masked classification are unsharpened, with the two other protomers removed for clarity. Colored classes were used for further model refinement (blue: OFS_in_; green: OFS_mid_; purple: OFS_out_, D390 down; wheat: OFS_out_, D390 up). All maps are contoured a σ of 8.

**Figure 4 – Supplementary Table 1.** Model refinement and validation statistics for Dataset B.

|  | P-Glt<sub>Ph</sub> OFS | P-Glt<sub>Ph</sub> OFS<sub>out</sub> | P-Glt<sub>Ph</sub> OFS<sub>out</sub> | P-Glt<sub>Ph</sub> OFS<sub>mid</sub> | P-Glt<sub>Ph</sub> OFS<sub>in</sub> |
| --- | --- | --- | --- | --- | --- |
| D390 state |  | down | up | down | down |
| **Cryo-EM acquisition and processing** |  |  |  |  |  |
| EMDataBank ID | EMD-24405 | EMD-24405 | EMD-24405 | EMD-24405 | EMD-24405 |
| Symmetry imposed | C3 | C1 | C1 | C1 | C1 |
| Particle classification | trimers | protomers | protomers | protomers | protomers |
| Initial particles | 9,234,148 | 1,620,456 | 1,620,456 | 1,620,456 | 1,620,456 |
| Final particles | 503,427 | 78,248 | 80,207 | 243,208 | 142,738 |
| Resolution (masked FSC = 0.143, Å) | 2.20 | 2.56 | 2.61 | 2.37 | 2.40 |
| Density modified CC (0.5, Å) | 2.20 | 2.56 | 2.59 | 2.33 | 2.39 |
| **Model Refinement** |  |  |  |  |  |
| PDB ID | 7RCP |  |  |  |  |
| Model resolution (FSC = 0.50/0.143 Å) | 2.23 / 2.01 | 2.60 / 2.08 | 2.66 / 2.04 | 2.36 / 1.92 | 2.42 / 2.00 |
| **Model composition** |  |  |  |  |  |
| Non-hydrogen atoms | 9339 | 9294 | 9294 | 9306 | 9307 |
| Protein residues | 1248 | 1246 | 1246 | 1246 | 1245 |
| Ligands | 6 | 6 | 6 | 6 | 6 |
| Waters | 57 | 29 | 31 | 41 | 49 |
| **R.m.s. deviations** |  |  |  |  |  |
| Bond lengths (Å) | 0.005 | 0.003 | 0.003 | 0.004 | 0.003 |
| Bond angles (°) | 0.620 | 0.494 | 0.488 | 0.618 | 0.550 |
| **Validation** |  |  |  |  |  |
| MolProbity score | 1.18 | 1.26 | 1.38 | 1.07 | 1.15 |
| Clash score | 3.88 | 5.00 | 5.52 | 2.79 | 3.58 |
| Poor rotamers (%) | 0 | 0 | 0 | 0 | 0 |
| **Ramachandran plot** |  |  |  |  |  |
| Favored (%) | 98.79 | 98.54 | 98.62 | 98.62 | 98.78 |
| Allowed (%) | 1.21 | 1.26 | 1.38 | 1.38 | 1.22 |
| Disallowed (%) | 0 | 0 | 0 | 0 | 0 |

**Figure 4 - Supplementary Figure 2.**
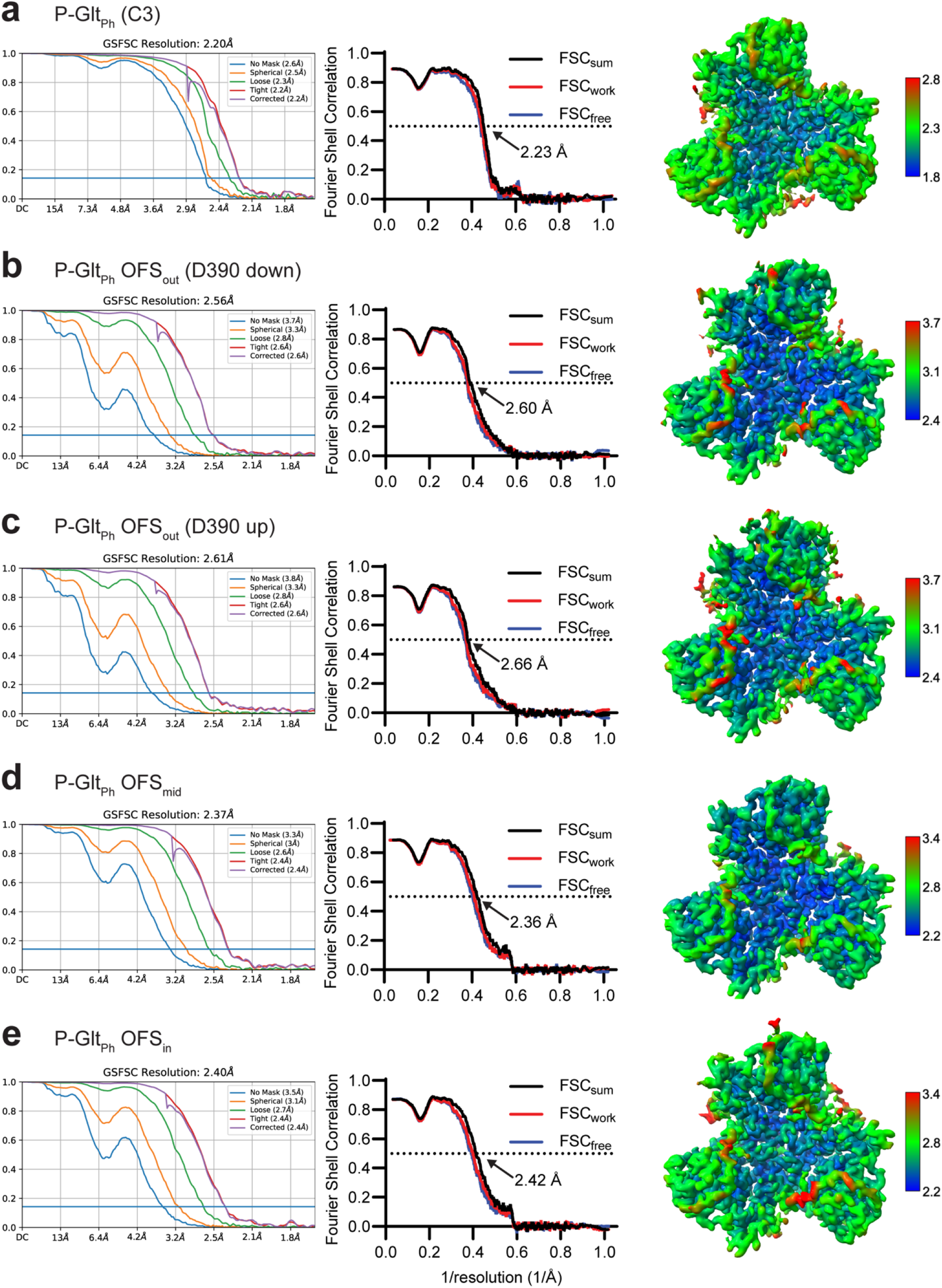
Validation statistics for models from Dataset B. (a) OFS-C3; (b) OFS_out_, D390 down; (c) OFS_out_, D390 up; (d) OFS_mid_; (e) OFS_in_. From left to right, map FSC from NU-refinement in cryoSPARC, model-to-data validation in Phenix of the trimer, and local resolution estimation. All maps are contoured at 8 σ except (A), which is contoured at 10 σ. The top protomer in (B-E) is the subject of focused classification and model refinement.

**Figure 4 – Supplementary Figure 3.**
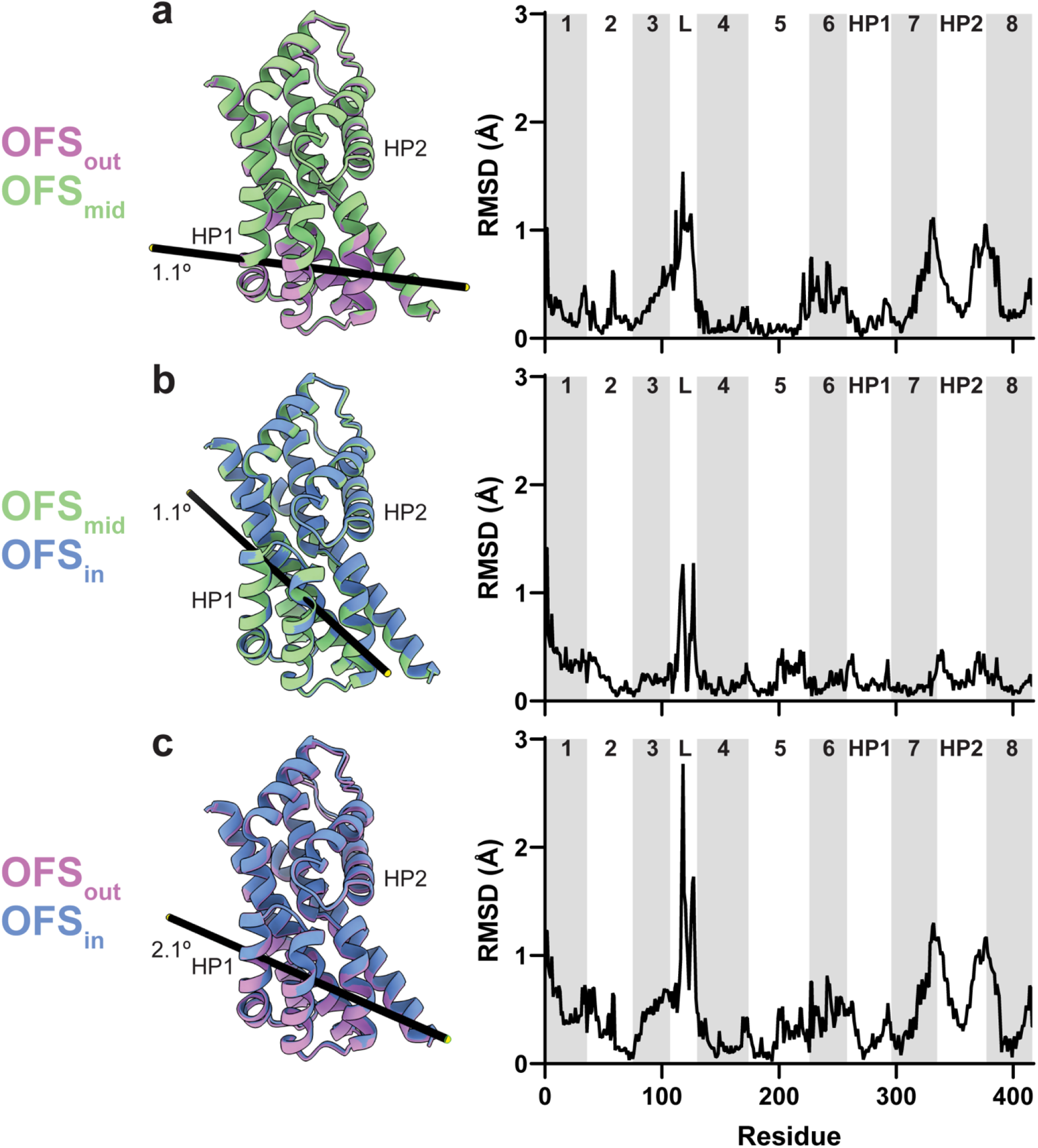
Protomer tilts in the outward-facing state. Trimers were superimposed on the trimerization regions (residues 150-195) of all three protomers. Comparison of the tilts for (a) OFS_out_/OFS_mid_; (b) OFS_mid_/OFS_in_; and (c) OFS_out_/OFS_in_. OFS_out_, OFS_mid_, and OFS_in_ are purple, green, and blue, respectively. Though parts of the scaffold domain also move (see Figure 5 – Movie 1), only transport domains of the classified protomers are shown for clarity. Black bars represent tilt axes and angles calculated using the ‘align’ command in ChimeraX. The corresponding per-residue Cα RMSDs are on the right. Transmembrane domains are labeled and alternatively shaded. ‘L’ denotes the flexible loop between TMs 3 and 4.

**Figure 4 – Supplementary Figure 4.**
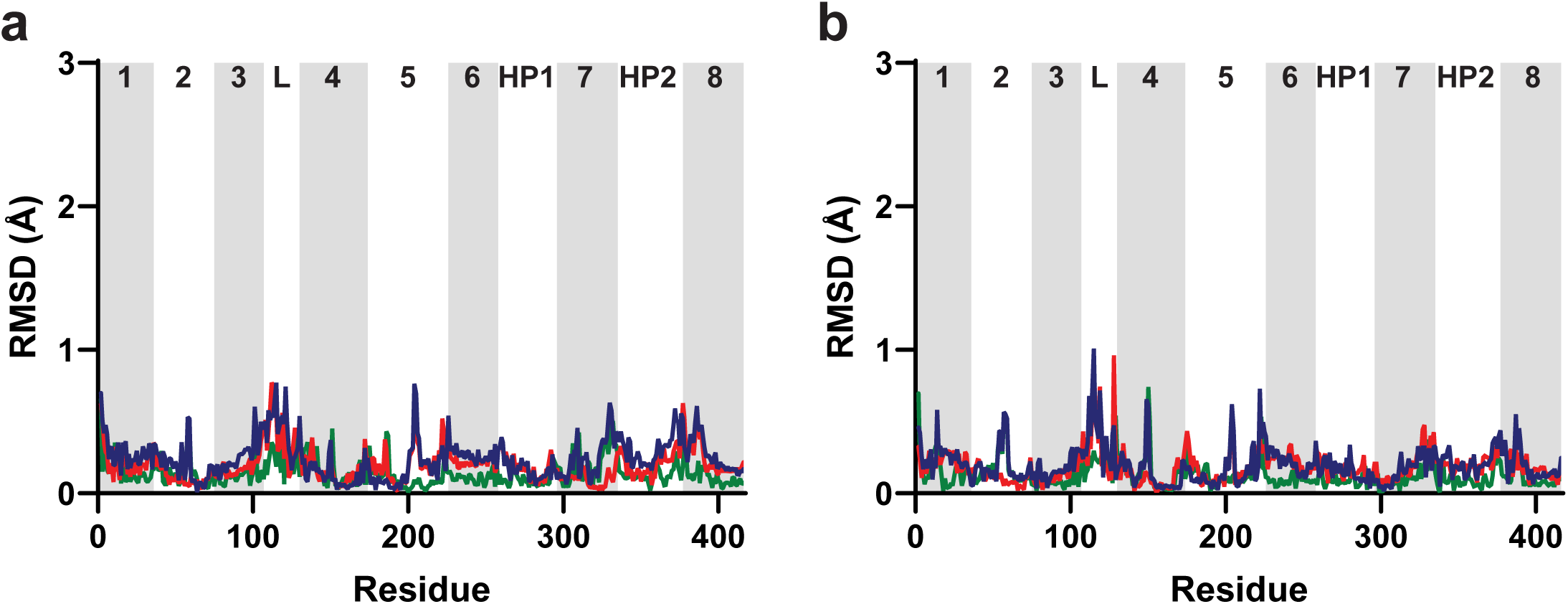
Protomers adjacent to chain A do not display concerted movements. Trimers were aligned along the trimerization regions of all three protomers (residues 150-195). Per-residue Cα RMSDs of (a) chain B, and (b) chain C. Lines are (green) OFS_out_/OFS_mid_; (red) OFS_mid_/OF_Sin_; and (blue) OFS_out_/OFS_in_. Transmembrane domains are labeled and alternatively shaded. ‘? indicates the disordered loop between domains 3 and 4.

**Figure 4 – Supplementary Table 2.**
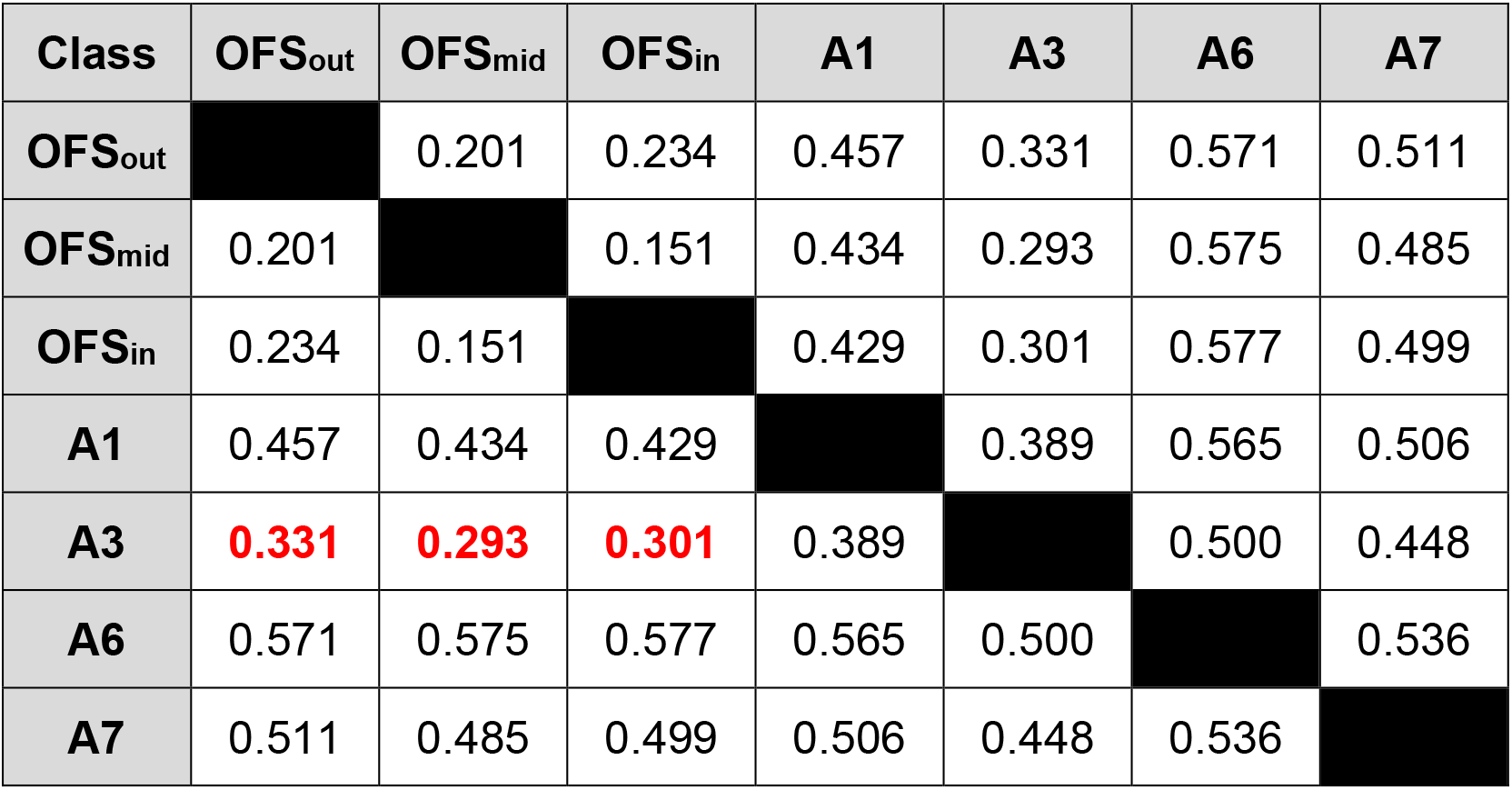
Comparison between structures from Dataset A and tilt states from Dataset B. Cα RMSDs were determined using ‘matchmaker’ implemented in ChimeraX using default parameters. Residues 12-106, 130-416 were used for structural alignment to include only well-structured, well-resolved regions. Lowest RMSD values between classes from Dataset A and Dataset B are red/bolded.

**Figure 4 – Supplementary Figure 5.**
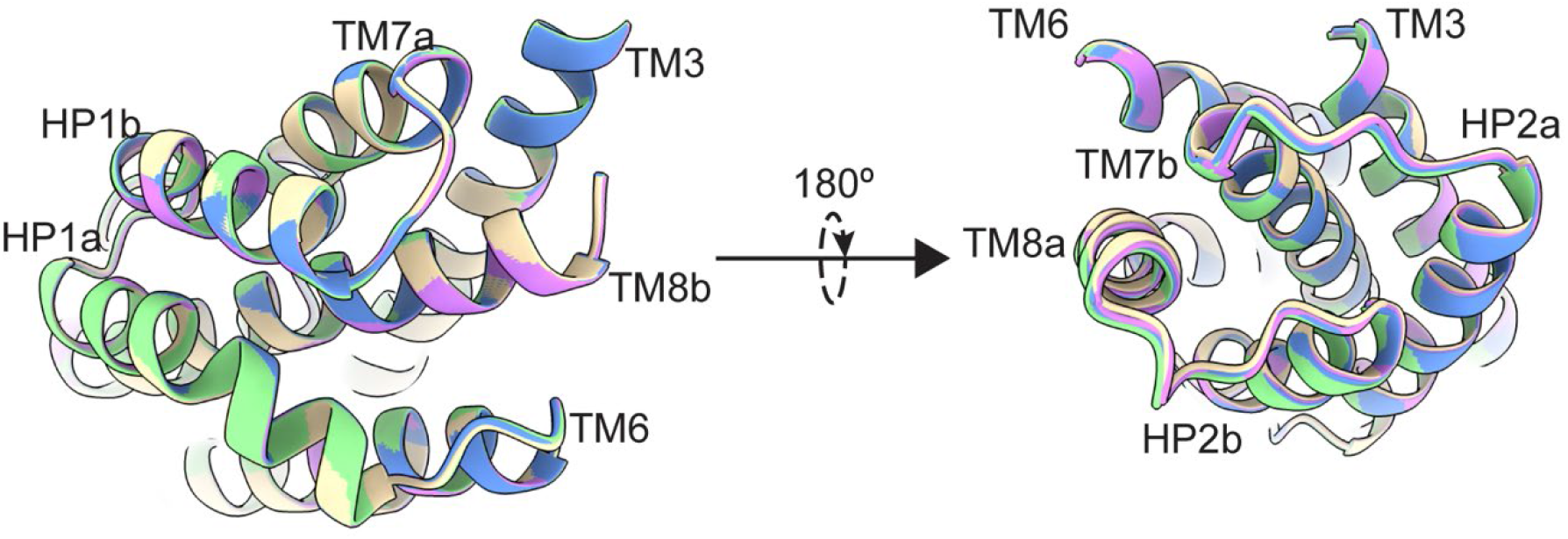
P-Glt_Ph_ at equilibrium does not have mobility in extracellular helices. Transport domains were superimposed on HP1 and TM7a (residues 258-309). Superimposition of transport domains from Dataset B. OFS_out_ (D390 down) is purple, OFS_out_ (D390 up) wheat, OFS_mid_ green, and OFS_in_ blue. The views are from the intracellular (left) and extracellular (right) sides of the transport domain.

**Figure 5 – Supplementary Table 1.** Correlation of tilt states and D390 rotamers from Dataset B processing.

| Class<br>(Round 1) | Tilt state | D390 rotamer | Class<br>(Round 2) | Tilt state | D390 rotamer |
| --- | --- | --- | --- | --- | --- |
| B1-1 | OFS<sub>out</sub> | intermediate | B2-1 | OFS<sub>out</sub> | down |
| B1-2 | OFS<sub>mid</sub> | unresolved | B2-2 | OFS<sub>mid</sub> | unresolved |
| B1-3 | OFS<sub>mid</sub> | down | B2-3 | OFS<sub>mid</sub> | down |
| B1-4 | OFS<sub>in</sub> | down | B2-4 | OFS<sub>out</sub> | up |
| **B1-5** | **OFS<sub>out</sub>** | **ambiguous** | B2-5 | OFS<sub>mid</sub> | unresolved |
| B1-6 | OFS<sub>mid</sub> | down | B2-6 | OFS<sub>mid</sub> | down |
| **B1-7** | **OFS<sub>mid</sub>** | **ambiguous** | B2-7 | OFS<sub>in</sub> | down |
| B1-8 | OFS<sub>mid</sub> | down | B2-8 | OFS<sub>mid</sub> | down |

